# A primary sensory cortical interareal feedforward inhibitory circuit for tacto-visual integration

**DOI:** 10.1101/2022.11.04.515161

**Authors:** Simon Weiler, Vahid Rahmati, Marcel Isstas, Johann Wutke, Andreas Walter Stark, Christian Franke, Christian Geis, Otto W. Witte, Mark Hübener, Jürgen Bolz, Troy W. Margrie, Knut Holthoff, Manuel Teichert

## Abstract

Tactile sensation and vision are often both utilized for the exploration of objects that are within reach though it is not known whether or how these two distinct sensory systems might combine such information. Here in mice we find that stimulation of the contralateral whisker array suppresses visually evoked activity in a subarea of primary visual cortex (VISp) whose visual space covers the whisker search space. This is mediated by local fast spiking interneurons that receive a direct cortico-cortical input predominantly from layer 6 of the primary somatosensory barrel cortex (SSp-bfd). These data demonstrate functional convergence within and between two primary sensory cortical areas for multisensory object detection and recognition.

## 1 Introduction

In everyday life, multiple types of sensory input arriving via distinct sensory modalities are simultaneously acquired to create a coherent and unified representation of the external world (Ernst and Banks, 2002; Gielen et al., 1983; Gleiss and Kayser, 2014; Gotz et al., 2017). The ability to rapidly and correctly recognize an object in the peripersonal space (Rizzolatti et al., 1981), (i.e. within reachable proximity), crucially relies on the orchestration of the tactile sensation and vision (Ernst and Banks, 2002). In rodents, both whisker based tactile sensation as well as vision are optimally combined to accurately evaluate the biological significance of nearby objects touched and seen simultaneously (Nikbakht et al., 2018; Shang et al., 2019). For instance, the performance of rats in judging the orientation of a solid object in close proximity dramatically increases when whiskers and vision work in concert (Nikbakht et al., 2018). Moreover, the interaction of these modalities is critically involved in prey capture-behavior in mice (Shang et al., 2019). Importantly, given that rodent’s whiskers are located in front of, or centered about their eyes (Huet and Hartmann, 2014), it is likely that both modalities operate within the same external space during object exploration. Thus, whisker-mediated tactile sensation and vision are deeply bound at the behavioral level and seemingly at the level of external sensory space.

In this study, we have used a combination of stereo photogrammetry for 3-dimensional (3D) reconstruction of the whisker array, intrinsic signal imaging, brain-wide viral retrograde and anterograde transsynaptic tracing followed by serial two-photon tomography and deep-learning based 3D detection of labeled cells, electrophysiology, optogenetics and mathematical network modeling to explore the possibility of tacto-visual convergence in the external proximity space and within the circuitry of the mouse VISp. We find that the search space of whiskers is highly associated with the visual space covered by VISp. Strikingly, this spatial multisensory convergence is precisely reflected in a subarea within VISp. Here, the anatomical location of postsynaptic excitatory neurons receiving direct cortico-cortical input from SSp-bfd, corresponds to the area in the visual space that overlaps with the external whisker search space. We further find that whisker stimulation has a powerful regulatory influence on VISp such that it cross-modally suppresses visually driven responses via fast-spiking interneuron mediated feedforward inhibition in layer 2/3. Our data reveal a specific anatomical and functional tacto-visual convergence at the level of VISp, highlighting the role of primary sensory areas in multisensory integration.

## 2 Results

### Mouse whiskers are prominently located in the visual space covered by VISp

As mouse’s whiskers are located in front of their eyes, we first aimed to explore to what extent they are associated with the external visual space covered by VISp. Because the 3D morphology of the mouse whisker array is unknown, we generated a morphologically accurate 3D model of this array based on stereo photogrammetry (Heist et al., 2018) data and aligned it with a realistic 3D model of the mouse head, including the eyes (Bolanos et al., 2021) (**Figures 1A, 1B, S1A-S1D**). Onto this model we then constructed the 3D visual space covered by VISp (Zhuang et al., 2017) (**Figure 1C**). Interestingly, already in this static model with whiskers and eyes in their intermediate positions, both whiskers and visual space show a market spatial overlap (**Video 1**, **Figure 1C**).

**Figure 1:**
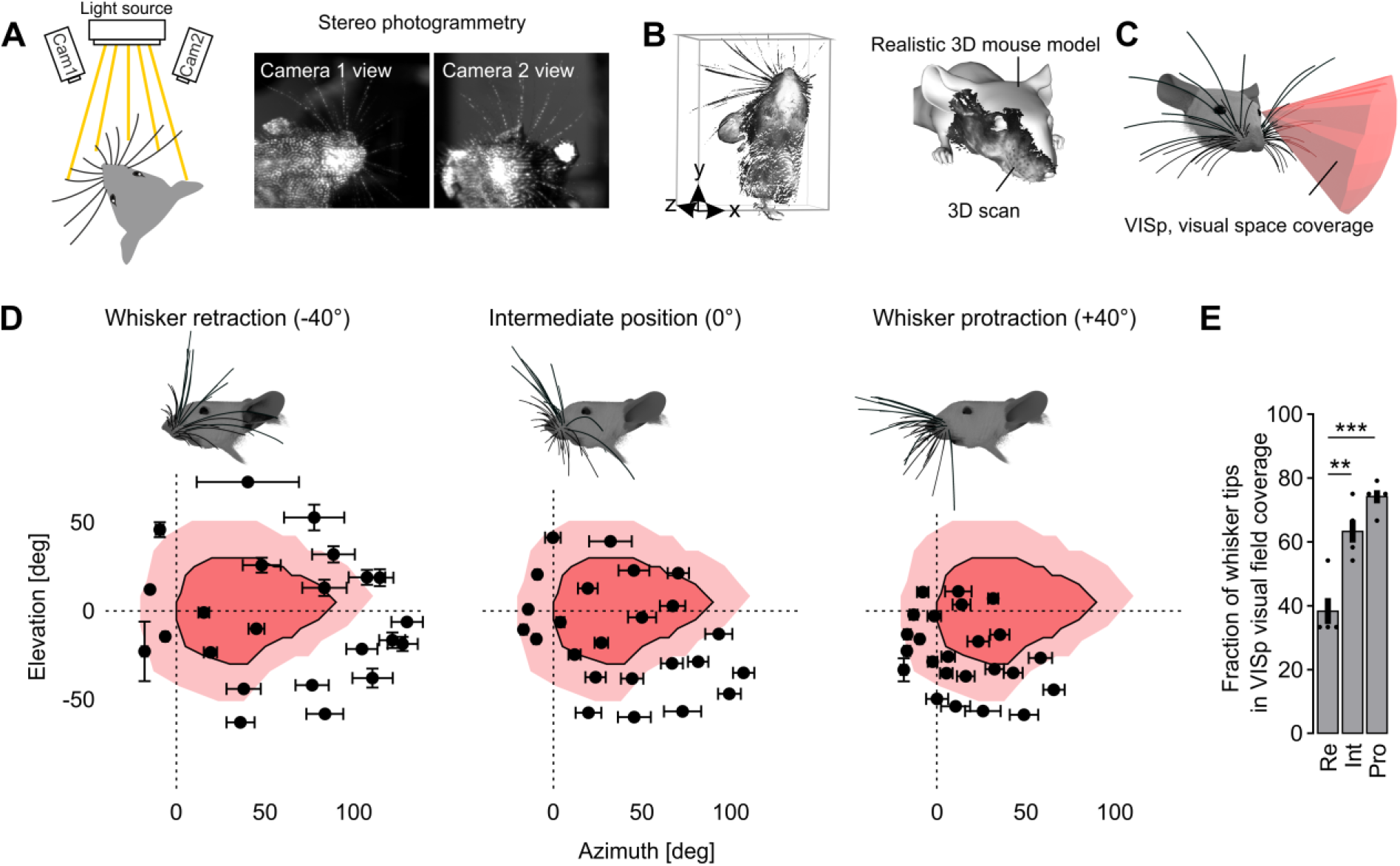
Tacto-visual overlap in the mouse proximity space. (A) The mouse head including the whisker array was illuminated with structured light patterns and stereo images were taken by two cameras. Detection of corresponding point pairs then allowed 3D-reconstruction via triangulation. (B) Left, Representative 3D point cloud of the mouse head with whiskers obtained after 3D-reconstruction. Right, Obtained 3D point clouds were aligned to an existing realistic 3D mouse model from (Bolanos et al., 2021). (C) 3D reconstructed and morphologically accurate model of the mouse head including eyes and whiskers (constructed in blender, see methods). Additionally, the 3D visual space covered by VISp originating from the left eye was constructed according to (Zhuang et al., 2017). (D) Mapping of tacto-visual overlap along azimuth and elevation. Dark pink area with surrounding solid black line: Coverage map of visual space by VISp (Zhuang et al., 2017). Bright pink area: Coverage map of visual space covered by VISp and mapped by eye movements in a ± 20 degree range. Centroids ± s.e.m. represent mean whisker tip positions under simulated retraction, intermediate and protraction conditions (n=5 mice). Mouse heads with whiskers display examples for whisker retraction, their intermediate position and whisker protraction (−40°, 0°, +40°). (E) Fraction of whisker tips located within the visual space covered by VISp under eye movement conditions during whisker retraction (Re), intermediate position (Int) and protraction (Pro). Black circles indicate data points of individual mice (Re vs. Int, p=0.0012; Re vs. Pro, p=0.0047; paired t-tests followed by Bonferroni correction) and bars indicate the mean fraction of whisker tips on total number of whiskers (the 24 large whiskers) ± s.e.m. **p<0.01, ***p<0.01

Mice typically gather sensory information by actively moving their sensory organs. More specifically, whiskers are rhythmically moved backward (retraction) and forward (protraction) during environmental exploration (Petersen, 2014). Additionally, mice move their eyes within an average ± 20 degree range around their central position (Meyer et al., 2020; Michaiel et al., 2020). Consequently, this shifts the visual space covered by VISp with respect to the location of the whisker-tips. To investigate the dynamic association of the actively scanned whisker and visual space, we simulated both whisker and eye movements (**Video 2**, **Figures S1H, S1I**, see methods). For quantification, we determined the average elevation and azimuth coordinates of the tip of each whisker on the left side of the snout, under retraction (−40°), intermediate (0°) and protraction (+40°) conditions in a left eye-centered spherical coordinate system (**Figures S1E-S1G**).

In all these conditions a substantial percentage of whisker-tips was present within the visual space (**Figure 1D**). Remarkably, while over the course of whisker protraction – a movement often associated with object exploration (Sofroniew and Svoboda, 2015) - the spread of the whisker array decreased (see also (Grant et al., 2009)), the number of whisker tips within visual space significantly increased from ~40% to ~80% (**Figures 1D, 1E**). Thereby, whisker tips accumulated in the lower, nasal visual space (**Figure 1D**). This implies, that mice can actively bring their tactile and (lower nasal) visual space into registration to operate within the same coordinate system. Thus, our data suggest that mice usually sense tactile and visual cues in proximity space in a spatially coherent fashion.

### Whisker stimulation suppresses visually driven activity in VISp *in vivo*

Having found that whiskers are prominently located in visual space, we wondered whether tactile sensation affects visual processing in VISp. Therefore, we explored the functional effects of tactile whisker stimulation on VISp activity using periodic intrinsic signal imaging (Kalatsky and Stryker, 2003). We measured visually driven VISp responses in restrained mice in the absence and presence of simultaneous whisker stimulation. The visual stimulus (v) was a vertically moving horizontal light bar displayed on a monitor in the nasal visual field of the left eye while whisker stimulation (w) was achieved by a vibrating metal pole moving continuously through the left whisker array, row by row. Independent stimulations evoked robust cortical activity and provided topographic maps of VISp and SSp-bfd, respectively (**Figures S2A, S2B**).

Remarkably, concurrent presentation of these stimuli significantly reduced the amplitude of visually driven VISp activity (**Figures 2A-2C**), suggesting a cross-modal modulation of VISp responses by tactile stimuli. Conversely, SSp-bfd responses remained unaffected by multisensory stimulation (**Figures S2C-S2E**), indicating an asymmetrical cross-modal effect. Next, concurrently with the visual stimulus we aimed to stimulate all whiskers on one side simultaneously. For this, we used air puffs generated by a picospritzer (**Figure 2D**). Interestingly, this stimulation led to an even stronger attenuation of visually elicited VISp responses (**Figures 2D’, S2K**), whereas ipsilateral whisker stimulation had no effect (**Figure S2J**).

**Figure 2:**
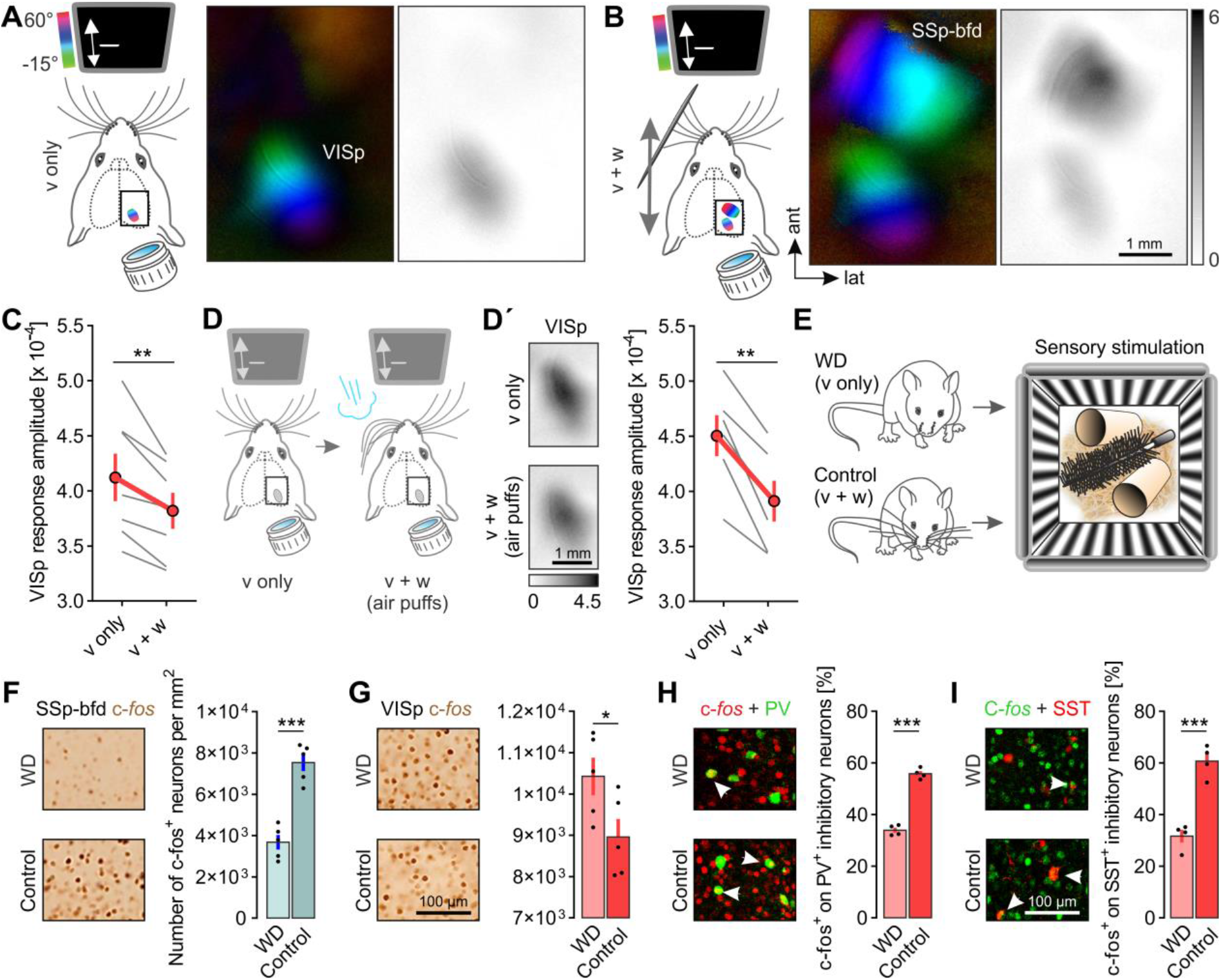
Whisker stimulation suppresses visually driven activity in VISp in vivo. (A, B) Schematics of unimodal visual (v only) and bimodal visual and whisker (v+w) stimulation procedures together with topographic and grey-scaled amplitude maps of VISp and SSp-bfd of one exemplary mouse. Darker amplitude maps indicate higher sensory evoked cortical activity. For bimodal stimulation both stimuli were temporally synchronized and spatially aligned. (C) Quantification of VISp response amplitudes under v only and v+w conditions (n=7, p=0.0067; paired t-test). (D) Schematic of the unimodal visual (v only) stimulation procedure and the bimodal visual and air puff induced whisker stimulation (v+w (air puffs)) procedure. (D’) Left, Grey-scaled amplitude maps of VISp obtained after unimodal and bimodal stimulation. Right, Quantification of VISp response amplitudes under v only and v+w conditions (n=6, p=0017; paired t-test). (E) Schematic of procedures preceding c-fos immunohistochemistry. Awake control mice and whisker deprived (WD) mice were placed in an environment for multisensory stimulation. The chamber consisted of four monitors displaying moving sine-wave gratings for visual stimulation. The bottom of the chamber was equipped with multiple densely arranged obstacles such as nesting material, paper roles and brushes enforcing simultaneous whisker stimulation during voluntary locomotion. (F) Left, Representative images of c-fos expression in SSp-bfd in WD and control mice after their exposure to the enriched environment. Right, quantified density of c-fos labeled neurons in SSp-bfd (n=5 mice per group, p=0.0001; unpaired t-test). (G) Left, representative images of c-fos expression in VISp in WD and control mice after their exposal to the same environment. Right, quantified density of c-fos labeled neurons in VISp (n=5 mice per group, p=0.0490, unpaired t-test). (H, I) Left, representative images of double immunostainings against c-fos and either PV or SST positive interneurons in VISp in WD and control mice after their exposure to the same environment. Right, quantified percentage of PV or SOM positive interneurons double labeled with c-fos on the total number of PV or SOM positive interneurons (n=4 mice per group, p=0.0000, p=0.0000; unpaired t-test). In C,D: Grey lines connect measurements of individual mice and red lines represent means ± s.e.m. In F-I: Black circles indicate measurements of individual mice and bars represents means ± s.e.m. *p<0.05, **p<0.01, ***p<0.01

We performed several control experiments to check for possible artifacts: (1) In whisker deprived (WD) mice, presenting the metal pole during visual stimulation did not lead to visual disturbances (**Figures S2F, S2F’**). (2) Likewise, air puffs also did not alter visually evoked activity in VISp in WD mice, indicating that sounds associated with the puffs did not contribute to the effect observed (**Figures S2G, S2G’**). (3) After eliminating the afferent input from the whiskers by cutting the infraorbital nerve (ION), whisker stimulation by air puffs had no effect on visual responses in VISp anymore, in contrast to sham surgery conditions (**Figures S2H, S2I**), indicating that that whiskers moving through visual space do not suppress VISp activity. Collectively, our data indicate a unihemispheric tacto-visual convergence at the level of VISp whereby tactile inputs act to globally suppress visually driven responses.

To investigate the effects of whisker stimulation on VISp activity in freely moving animals, we next used functional neuroanatomy to examine unrestrained mice after they were exposed to an enriched environment (**Figure 2E**, right). Two groups of mice were used, a control group with intact whiskers and an experimental WD group (**Figure 2E**, left). Thus, voluntary locomotion through the provided environment led to bimodal visual and whisker stimulation (v+w) in control, but visual stimulation alone (v only) in WD mice. We quantified neuronal activation in VISp and SSp-bfd using the expression of the immediate-early gene c-fos. As expected, the number of c-fos positive neurons in SSp-bfd was markedly higher in control compared to WD mice (**Figure 2F**). Strikingly, the opposite effect was observed in VISp, where we detected significantly less c-fos labeled neurons in control compared to WD mice (**Figure 2G**). Hence, we conclude that also in unrestraint mice visual responses in VISp are reduced by concurrent whisker stimulation. Because about 80% of all cortical neurons are excitatory, this effect can predominantly be attributed to a reduced responsiveness of these neurons in VISp. To address the contribution of inhibitory GABAergic neurons separately we specifically determined c-fos expression in parvalbumin (PV) and somatostatin (SST) positive cells. Both, PV and SST expressing inhibitory neurons showed significantly higher *c-fos* expression-levels in control mice (**Figures 2H, 2I**) indicating that whisker stimulation cross-modally drives local inhibitory circuits in VISp.

### Layer 6 excitatory neurons in SSp-bfd are the main source for direct projections to VISp

Next, we aimed to identify the pathway underlying tactile integration in VISp since the source of whisker-related inputs in VISp is unknown. To systematically identify neurons projecting to VISp, we utilized retrograde tracing using a self-engineered recombinant AAV variant, AAV-EF1a-H2B-EGFP, which leads to the expression of EGFP in the nuclei of projection neurons (nuclear retro-AAV). This virus was injected into different positions across the extent of VISp, whereby each mouse received one injection extending across all cortical layers (**Figures 3A, 3B, 3I**). Following brain-wide *ex vivo* two-photon tomography, retrogradely labeled neurons across the entire brain were counted using a deep learning based algorithm for 3D cell detection (Tyson et al., 2021) and assigned to brain areas of the Allen Mouse Brain Common Coordinate Framework (CCFv3) (Wang et al., 2020) (**Figure S3B**).

**Figure 3:**
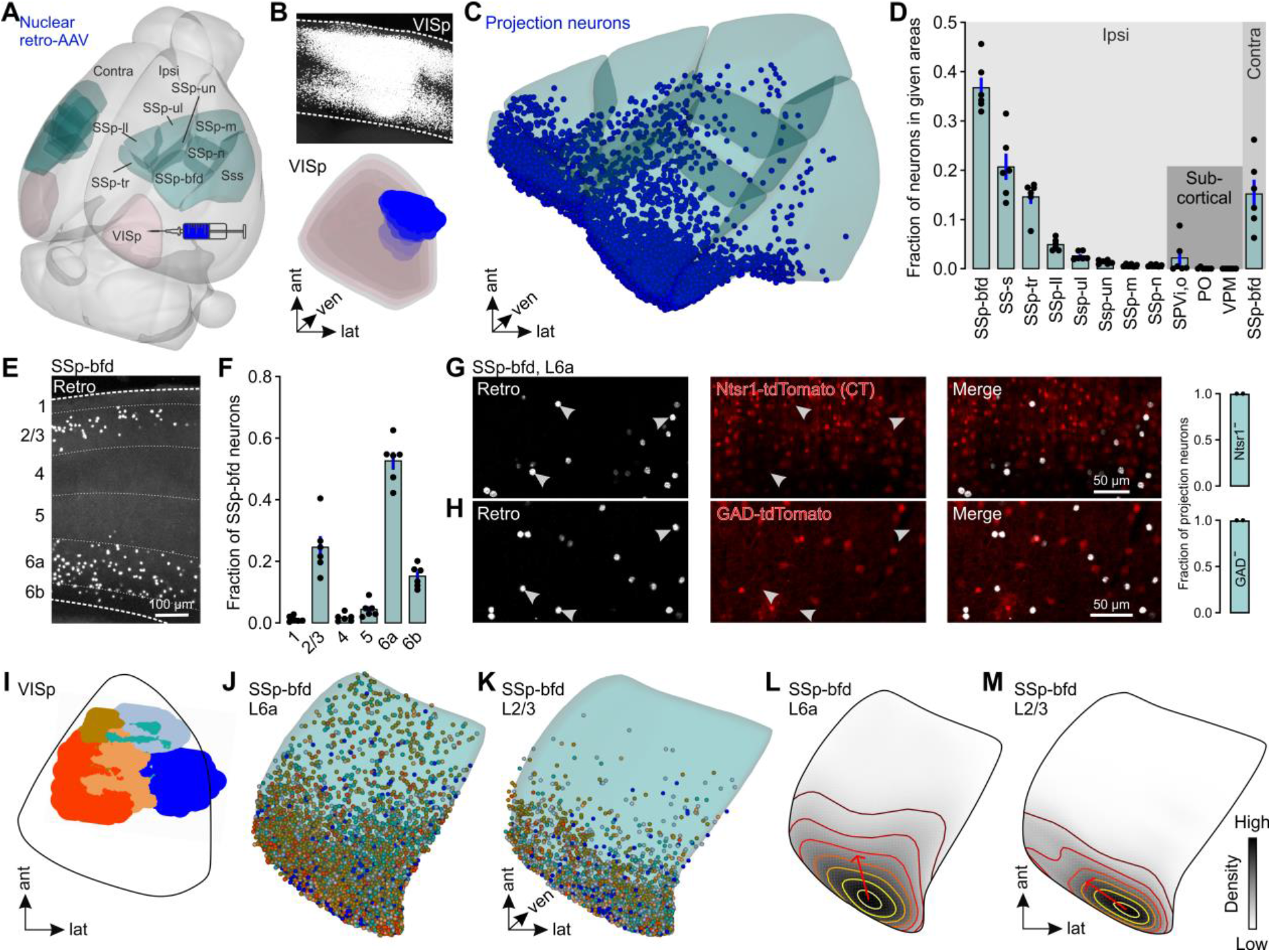
Excitatory cortico-cortical (CC) neurons in L6 in SSp-bfd are the main source for direct projections to VISp. (A) 3D rendered mouse brain showing the locations of the somatosensory cortical areas and VISp together with a schematic of the viral injection approach. (B) Top, coronal section showing a representative injection site in VISp. Bottom, 3D reconstruction of the same injection site warped into the 3D rendered space of VISp of the CCFv3 (Wang et al., 2020). (C) Visualization of detected projection neurons of the same mouse warped into the 3D rendered space of cortical somatosensory areas of the CCFv3. (D) Fraction of projection neurons in different cortical somatosensory areas and whisker-recipient subcortical areas (n=6). Black circles represent fractional cell counts of individual animals. Bars represent means ± s.e.m. (similar across all plots). (E) Representative coronal section showing projection neurons (white) in SSp-bfd. Numbers indicate cortical layers. (F) Fraction of projection neurons across different layers of SSp-bfd (n=6). (G, H) Coronal sections showing tdTomato expression and retrogradely labeled neurons in L6 in SSp-bfd of Ntsr1 and GAD-tdTomato mice (n=2 mice per group). (I) Reconstructed injection sites from 6 different mice warped to a horizontal projection of VISp. (J, K) Dorsal view to L6a and L2/3 of SSp-bfd. Detected projection neurons from the 6 different mice warped to L6a and L2/3 of SSp-bfd of the CCFv3. Colors relate to the injection sites in (I). (L, M) Average density of projection neurons in L6 and L2/3 of SSp-bfd (horizontal projection). The colors of the contour lines indicate cell density (yellow: high, dark brown: low). Closer distances between two contour lines reflect a steeper slope of density changes. The red arrows indicate the direction of the first principal component (PC) explaining the largest variance in the distribution of projection neurons (n=6).

We found that VISp receives projections from a large number of ipsilateral cortical and subcortical brain areas (**Figure S3A**). Importantly, when focusing on somatosensory brain areas, projection neurons were particularly abundant in the whisker-recipient SSp-bfd, while subcortical whisker-recipient areas only contained a negligible number of them (**Figures 3C, 3D, S3C**). Similar results were obtained when we injected another AAV-based retrograde tracer, rAAV2-retro.CAG.GFP that permits efficient access to cell bodies of projection neurons (cellular retro-AAV) (Tervo et al., 2016) (**Figure S3D**). Together, our data indicate that SSp-bfd is the major source for direct connections to VISp. Thus, this pathway is a promising candidate to directly mediate the functional effects of whisker stimulation on visually driven activity in VISp as observed above.

Within SSp-bfd and other subareas of SSp the dominant location of projection neurons was layer 6 (L6) followed by L2/3 (**Figures 3E, 3F**). Likewise, also in the primary auditory cortical area (AUDp), which has been shown to send direct functional connections to VISp (Ibrahim et al., 2016; Iurilli et al., 2012), L6 contained a substantial fraction of projection neurons as well, beside smaller fractions in L2/3 and 5 (**Figures S3G, S3H**). Importantly, L6 projection neurons in SSp-bfd were excitatory and non-overlapping with cortico-thalamic (CT) cells, the main cell type in L6 (Zhang and Deschenes, 1997), as revealed by virus injections in GAD-tdTomato or Ntsr1-tdTomato mice (Bortone et al., 2014), expressing tdTomato in GABAergic and CT neurons, respectively (**Figures 3G, 3H**). Projection neurons in L2/3 did also not co-express GAD-tdTomato (data not shown). Taken together, these data indicate that the location of projection neurons in L6 is a general feature of cross-modal cortico-cortical communication and that L6 cells projecting from SSp-bfd to VISp are cortico-cortical (CC) projection neurons.

We next asked whether there was any spatial organization of L6 and L2/3 projection neurons across SSp-bfd. Interestingly, independent of the different locations of the injection sites within VISp (**Figure 3I**), the highest density of projection neurons in both layers was observed in the posterior region of SSp-bfd, the part of SSp-bfd situated in close proximity to VISp (**Figures 3J-3M, S3J, S3L, S3M**). Projection neurons in both layers were not obviously organized topographically (**Figure S3K**).

### Projection neurons in SSp-bfd are located in the posterior barrel columns, which correspond to the most caudal whiskers

In rodents, SSp-bfd contains discrete clusters of neurons in L4, called “barrels” which are arranged somatotopically in an identical pattern as the whiskers on the snout (Van der Loos and Woolsey, 1973). Thereby, each whisker preferentially innervates one barrel and its cortical column (Petersen, 2019). Here, we investigated the association of neurons projecting to VISp with these barrel columns. For this, we first reconstructed the entire barrel field in layer 4 from an average autofluorescence image data set obtained by serial two-photon tomography of 1675 mouse brains (Wang et al., 2020), using the brainreg-segment software (Tyson et al., 2022) (**Figures 4A-4D**). Next, to generate a map of the barrel columns in L6 and L2/3 of SSp-bfd, the reconstructed barrel field was scaled into these layers.

**Figure 4:**
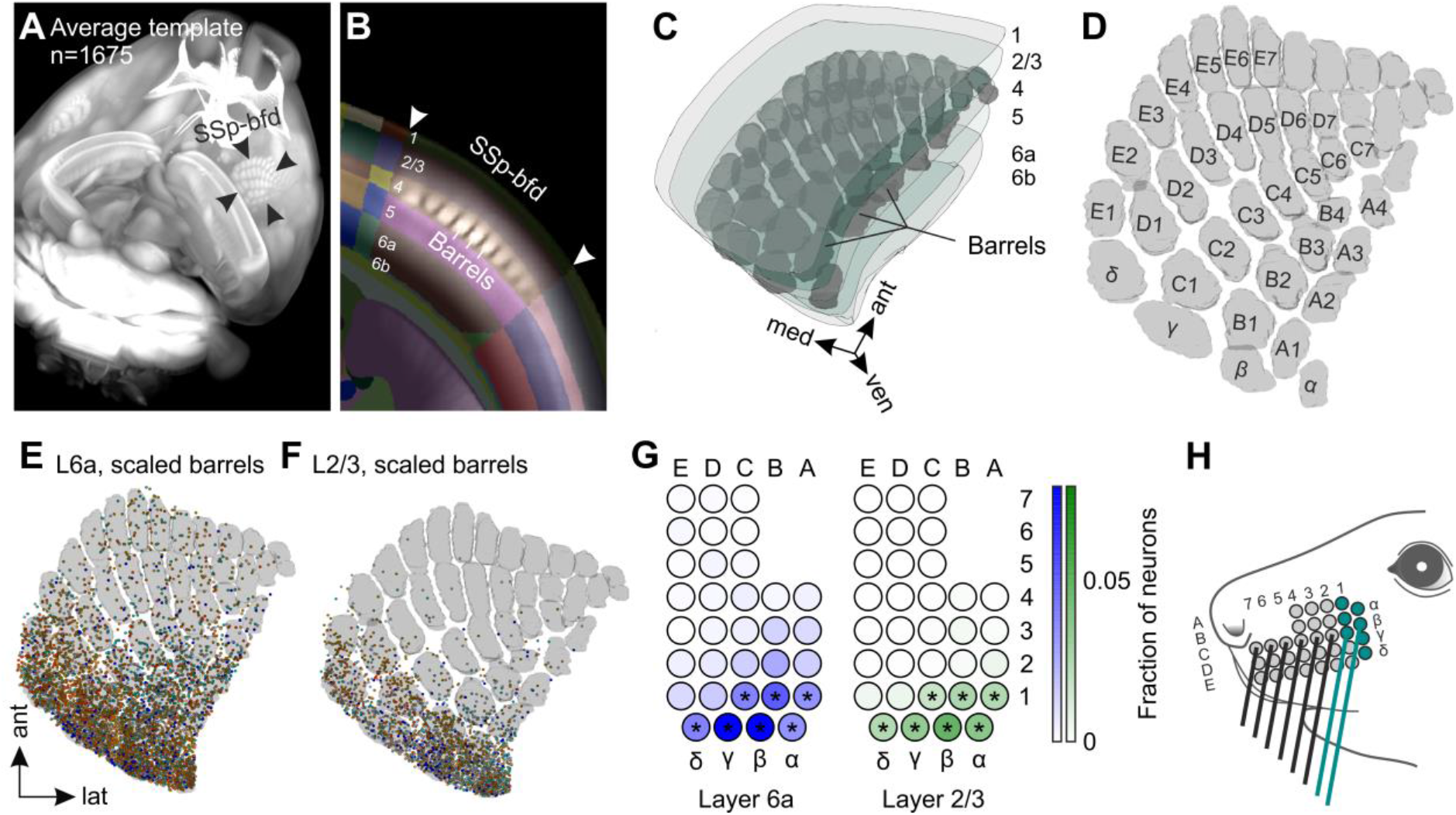
Projection neurons in caudal barrel columns innervated by caudal whiskers are the main source for direct projections to VISp. (A) Mouse brain reconstructed in 3D from an autofluorescence image data set obtained by serial two photon tomography of 1675 mouse brains (Wang et al., 2020). (B) The autofluorescence imaging data set was aligned to theCCFv3 (Wang et al., 2020) using the brainreg software (Tyson et al., 2022). Barrels became visible in L4 of SSp-bfd. Numbers indicate cortical layers. (C) Barrels in L4 were reconstructed in 3D using brainreg-segment (Tyson et al., 2022) and warped to the 3D rendered space of SSp-bfd of the CCFv3 (Wang et al., 2020). (D) Horizontal projection of the reconstructed barrels together with the standard nomenclature for rows (A-E) and arcs (1-7). (E, F) Overlay of projection neurons labeled by the 6 different injections sites in VISp with reconstructed barrel field scaled in L6 and L2/3 of SSp-bfd (horizontal projections). (G) Color-coded output maps showing relative average output strength of individual barrel columns in L6 and L2/3 in SSp-bfd (n=6). Asterisks indicate barrel columns containing a significant number of projection neurons as compared to uniformly distributed cell positions (*p<0.001). Values are normalized to the total number of neurons within all barrel columns in both L6a and L2/3. (H) Schematic of the mouse mystacial pad. Circles indicate whisker basepoints. Colored whiskers represent the ones predominantly innervating the barrel columns in SSp-bfd with strong significant projections to VISp.

**Figures 4E and 4F** illustrate an overlay of the barrel-columns with the projection neurons labeled by the different injection sites in VISp. Generally, projection neurons in both L6 and L2/3 were detected within the barrel columns and their separating septa. As expected, we identified specifically the posterior barrel columns to contain a significant amount of neurons, which was beyond chance levels, in both layers, compared to a modeled uniform distribution of projections neurons across SSp-bfd (**Figure 4G**). These posterior columns are preferentially innervated by the most caudal whiskers (**Figure 4H**). This suggests, that somatosensory information, particularly gathered in the space scanned by the caudal whiskers, plays a crucial role in multisensory tactile integration in VISp.

### Locations of postsynaptic neurons in VISp correspond to the lower lateral visual field

Given the observed anatomical projections from SSp-bfd to VISp, we next aimed to explore the precise location and spatial distribution of postsynaptic neurons in VISp. For this, we utilized AAV-meditated anterograde trans-synaptic tracing in which the injection of a virus containing Cre-recombinase (AAV2/1-hSyn-Cre) in the presynaptic neuronal population induces the conditional expression of a reporter gene in postsynaptic neurons (Zingg et al., 2017). We injected this virus into different positions spanning the extent of SSp-bfd in Ai14 mice (**Figures 5A, 5B**).

**Figure 5:**
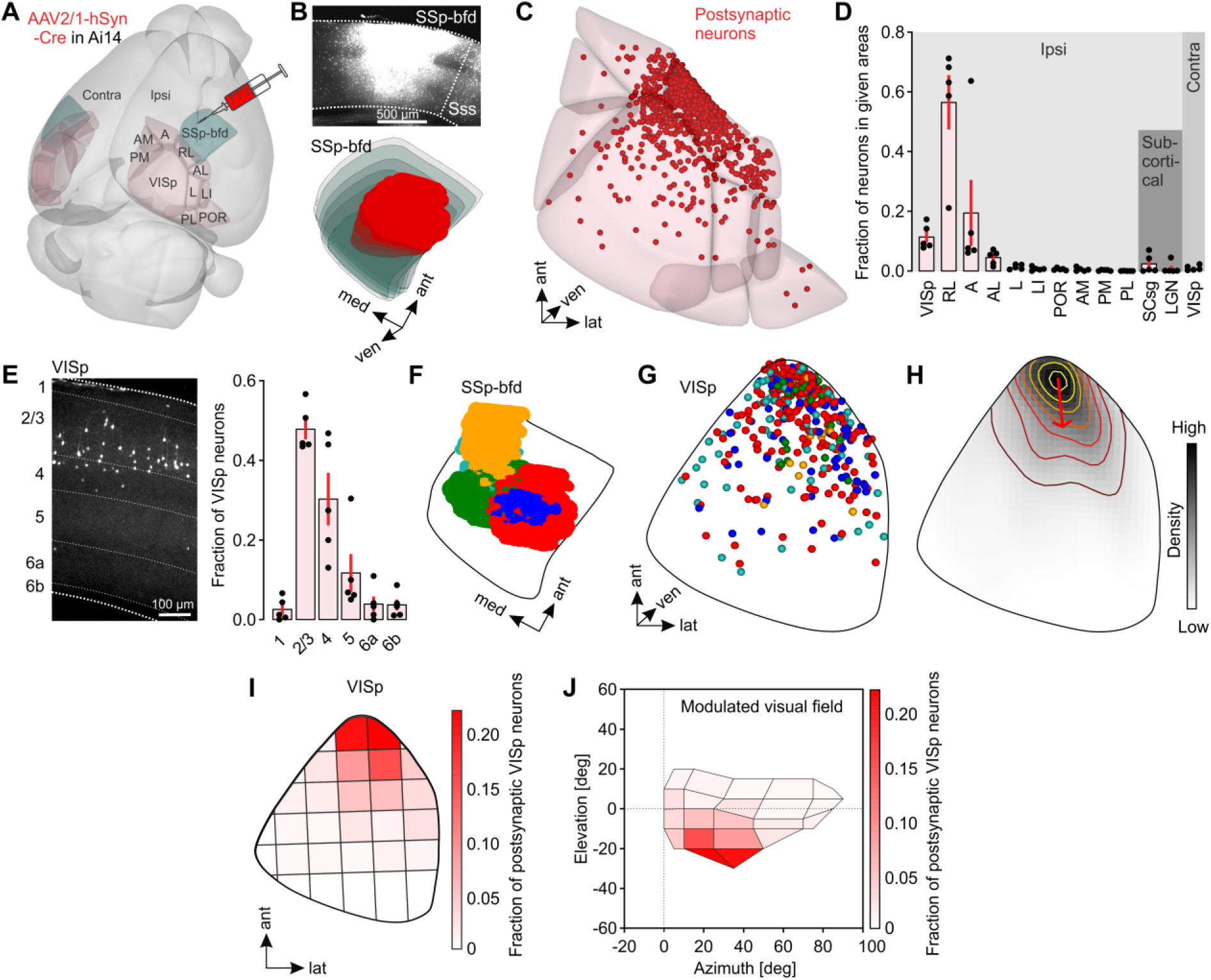
Target neurons in VISp are mainly located in L2/3 and their location in elevation and azimuth corresponds to the lower, nasal visual field. (A) 3D-rendered mouse brain showing the locations of SSp-bfd and visual cortex areas together with a schematic of the viral injection approach. (B) Top, coronal section showing a representative injection site in SSp-bfd. Bottom, 3D reconstruction of the same injection site warped into the 3D rendered space of SSp-bfd of the CCFv3 (Wang et al., 2020). (C) Visualization of detected postsynaptic neurons of one representative mouse warped to cortical somatosensory areas of the CCFv3 of visual cortex areas. (D) Fraction of postsynaptic neurons in different cortical and subcortical visual areas (n=5). Black circles represent fractional cell counts of individual animals. Bars represent means ± s.e.m. (similar across all plots). (E) Left, representative coronal image showing postsynaptic neurons (white) in VISp. Numbers indicate cortical layers. Right, fraction of postsynaptic neurons across different layers of VISp (n=5). (F) 3D-reconstructed injection sites from 5 different mice warped to a horizontal 2D projection of SSp-bfd. (G) Detected postsynaptic neurons from 5 different mice warped to VISp of the CCFv3 (horizontal projection). (H) Average density of projection neurons in VISp (horizontal projection, n=5). Contour lines indicate the slope of density changes and the red arrow indicates the first principal component explaining the largest variance in the distribution of postsynaptic neurons (n=5 mice). (I) Horizontal projection of VISp. Parcellation was performed based on PCA of the postsynaptic neuron distribution. The color-coded map shows the average fraction of postsynaptic neurons (n=5 mice) in each parcel. (J) Visual space covered by the detected postsynaptic neurons in VISp. Color-coded is the average fraction of postsynaptic neurons (n=5 mice) in each parcel of VISp.

We found postsynaptic neurons in multiple cortical and subcortical areas, known to be directly targeted by SSp-bfd (Petersen, 2019) (**Figures S4A, S4B**). Strikingly, labeled neurons were also abundant in visual cortical areas including higher-order visual areas (HVAs) and VISp (**Figures 5C, 5D, and S4D**). In contrast, we only detected a negligible number of postsynaptic neurons in subcortical visual areas (**Figure 5D**). In line with our observations made by retrograde tracing experiments, these data suggest that tactile integration in mouse visual cortex is substantially mediated by direct cortico-cortical connections originating in SSp-bfd.

We found that postsynaptic neurons in HVAs and VISp were preferentially located in L2/3 (**Figures 5E, S4D**), whereas postsynaptic neurons in AUDp were predominantly found in L5 and 6 (**Figures S4E-S4F**). This strongly suggests that L2/3 plays an important role for cross-modal integration of tactile inputs within visual areas. Our retrograde tracing results suggest that whisker-related tactile inputs in VISp originate in L6 and L2/3 of SSp-bdf. To investigate which cortical layers in VISp are targeted by these specific projection sources, we injected the anterograde tracer specifically into L6 or L2/3 in SSp-bfd (**Figures S4H, S4I**). Strikingly, the vast majority of postsynaptic neurons in VISp was found after L6 injections (**Figure S4I**, left). These neurons were predominantly located in L2/3 (and L5, **Figure S4I**, right) suggesting that the direct pathway from L6 in SSp-bfd to L2/3 in VISp is critically involved in mediating the effects of whisker stimulation on VISp responses.

Within VISp postsynaptic neurons were not obviously topographically arranged and had the highest density in the anterior part of VISp (**Figures 5G, 5H, S4J, S4K**), independent of the injection positions within SSp-bfd (**Figure 5F**). Accordingly, PCA-based parcellation of VISp revealed a steep gradient in the number of postsynaptic neurons in anterior-posterior direction (**Figures S4L, S4M**). These data indicate that SSp-bfd mediated tacto-visual convergence is restricted to a transitional zone in the anterior part of VISp located in close proximity to SSp-bfd.

Finally, we examined the association between the location of postsynaptic neurons and the functional spatial organization of VISp. In detail, VISp contains a continuous representation of the contralateral visual field (Drager, 1975; Wagor et al., 1980). The lower visual field is represented in the anterior part (elevation), and the nasal visual field innervates the lateral part of VISp (azimuth) (Drager, 1975; Garrett et al., 2014; Wagor et al., 1980; Zhuang et al., 2017). To investigate the association of postsynaptic neurons with the visual field representations, we first parceled VISp into 31 subareas, based on non-arbitrary sectioning along the first and second principal component (PC) of the postsynaptic neuron distribution, and determined the average fraction of labeled neurons in each of these subareas (**Figure 5I**). Then, we assigned visual space coordinates (Zhuang et al., 2017) (**Figures S4N, S4O**) to each subarea to estimate its visual space coverage. Remarkably, we found that subareas with high cell counts represent the lower, nasal visual space (**Figure 5J**), reminiscent of the search space of whiskers under protraction conditions. This implies that visual signals predominantly from this part of visual space are suppressed by SSp-bfd inputs.

### SSp-bfd functionally targets VISp

Given the prominent cross-modal projections from SSp-bfd to VISp, we sought to delineate the functional strength and specificity underlying these anatomical connections. For this, we injected the SSp-bfd with AAV.CaMKIIa-hChR2 tagged with EYFP to express light sensitive cation channels in excitatory cells (**Figure 6A**). Additionally, we co-injected AAV.Syn.Cre to anterogradely label potential postsynaptic targets in VISp of Ai14 mice. With this approach, we observed both axonal fibers expressing ChR2 and anterogradely labelled cells with the highest density in the anterior part of VISp similar to our previous observations (see **Figures 5 and 6B**). We then used whole-cell patch clamp recordings to measure optically evoked postsynaptic currents or potentials (PSCs or PSPs) of L2/3 cells in acute brain slices of VISp.

**Figure 6:**
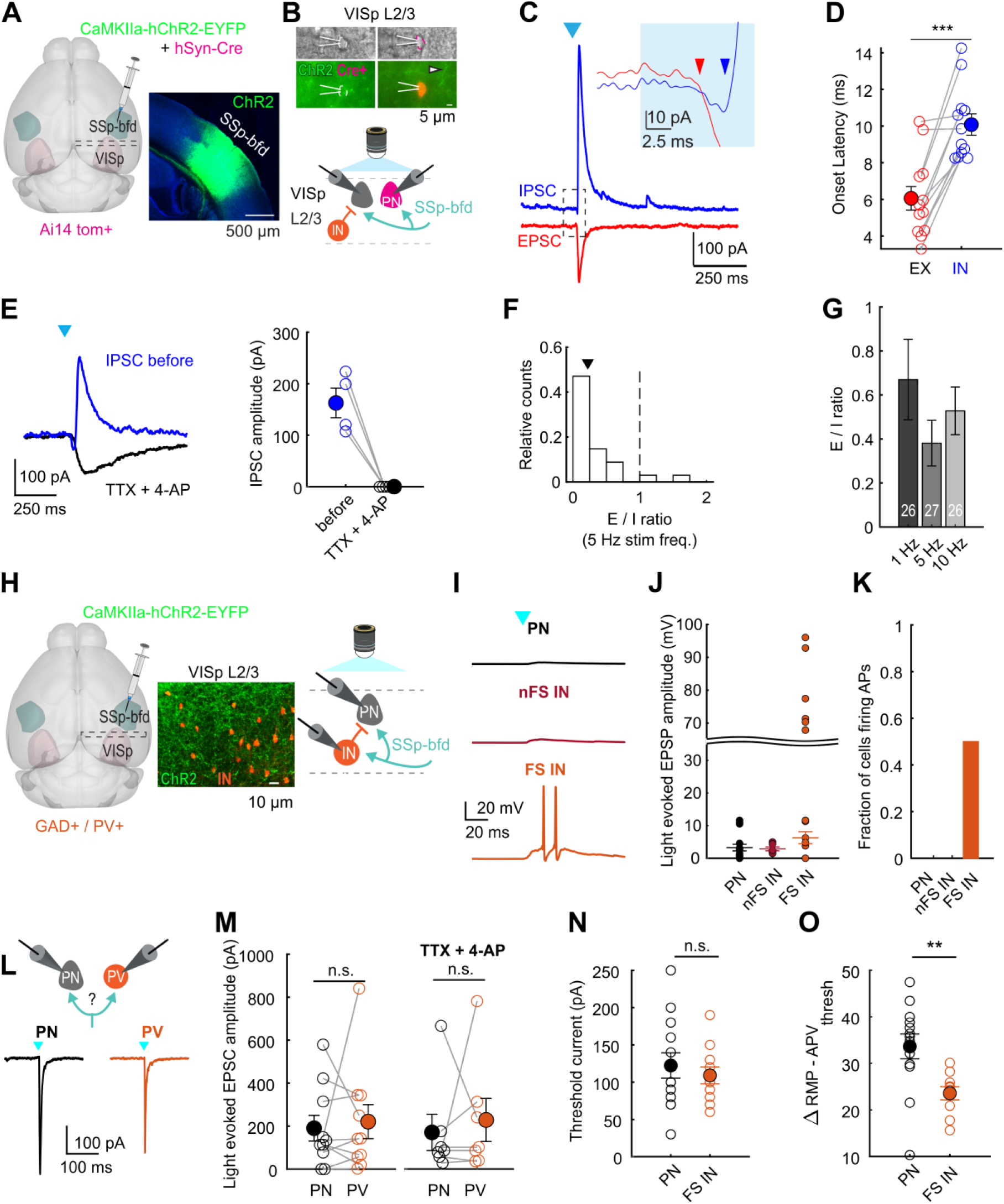
SSp-bfd mediated feedforward inhibition onto L2/3 neurons in VISp. (A) Schematic of viral injection approach (left). AAV.CaMKIIa-hChR2-EYFP and AAV.hSyn.Cre were co-injected across all layers in SSp-bfd in Ai14 mice. Right, Expression of AAV.CaMKIIa-hChR2 tagged with EYFP in SSp-bfd. (B) Top, epifluorescence images of Cre- and Cre+ L2/3 neurons in VISp during patch recording. Cre+ cells are anterogradely labelled cells (see also S5A and Methods). Arrow highlights axonal fibers expressing AAV.CaMKIIa-hChR2-EYFP (green). Bottom, Schematic of recording and photostimulation configuration. Neighboring Cre+ and Cre-cells were recorded in VISp while axonal fibers from SSp-bfd were activated by 472 nm light. (C) Representative example of light-evoked inhibitory and excitatory postsynaptic current (IPSC in blue and EPSC in red) in L2/3 pyramidal neuron (PN). Cell is clamped at 0 and −70 mV holding potentials, respectively. SSp-bfd fiber stimulation elicited a short-latency monosynaptic EPSC followed by a delayed IPSC. Blue arrowhead indicates light onset. Inset, enlarged view of the circled area, in which the two arrows indicate the onset of the EPSC and IPSC. (D) Onset latencies of light-evoked IPSCs were significantly longer than that of light-evoked EPSCs. Mean (filled circles) and individual data points (empty circles) are displayed. Lines connect peak EPSC and IPSC of the same cell (n=12 cells from 6 mice). Error bars are s.e.m., p<0.001, Wilcoxon signed rank. (E) Left, Representative IPSC recorded before (blue) and after infusion of TTX and 4-AP (black). Right, Comparison between peak amplitude of IPSCs before and after infusion of TTX and 4-AP. Mean and individual data points are displayed (n=4 cells from 4 mice). Error bars are s.e.m. (F) Distribution of E/I ratio across recorded cells (Cre+ and Cre-pooled, see S5B, S5C). Data are shown for 5 Hz light stimulation using peak response amplitudes for EPSCs and IPSCs (n=27 cells, from 7 mice). (G) Mean E/I ratio for different stimulation settings (1 Hz, 100ms long; 5Hz, 10ms; 10Hz, 10ms). Error bars are s.e.m. Cell numbers are indicated. (H) Injection scheme for GAD/PV animals. Injection approach like (A). Middle, axonal fibers expressing AAV.CaMKIIa-hChR2 tagged with EYFP (green) and PV+ interneurons expressing tdtomato (orange). Right, schematic of recording and photostimulation configuration. (I) Example light-evoked postsynaptic potentials (EPSPs) for L2/3 PNs, non-fast spiking and fast spiking interneurons (nFS IN, FS IN). Arrowhead indicates light onset. Note that FS INs fired action potentials upon light activation. (J) Light-evoked sub- and suprathreshold EPSPs for PNs, nFS INs and FS INs. Suprathreshold action potential firing is displayed above curved line. (K) Fraction of cells showing light-evoked action potential firing. (L) Top, Recording configuration to probe light-evoked responses in neighbouring L2/3 PNs and PVs (FS INs). Bottom, Representative example of light-evoked EPSCs in PN and FS IN. (M) Comparison of evoked peak EPSCs for PNs and FS INs without (left) or with TTX and 4-AP present (right). Mean (filled circles) and individual data points (empty circles) are displayed. Lines connect neighboring cells (<100 μm apart; n=11 cells and n=7 cells from 3 and 2 mice). Error bars are s.e.m. (p=0.85, p=0.65 Wilcoxon rank-sum). (N) Comparison of minimal amplitude of injected step current (Rheobase) that elicited action potential firing for PNs and FS INs (n=13 cells, n=11 cells from 6 mice, p=0.75 Wilcoxon rank-sum). (O) Comparison of membrane potential difference between resting membrane potential and spike threshold for PNs and FS INs (n=13 cells, n=10 cells from 6 mice, p<0.001, Wilcoxon rank-sum).

We first recorded from pyramidally-shaped tdtomato^+^ and neighbouring tdtomato^-^ neurons in L2/3 (PNs, less than 100 μm apart, **Figures 6B**, **S5A**). We observed both light-evoked excitatory and inhibitory PSCs in the same cells (**Figure 6C**). Given that the strength and latencies of light-evoked PSCs in tdtomato+ and tdtomato-negative cells were indistinguishable, we pooled these data together for the remaining analysis (**Figures S5B, S5C**). While the onset latencies of EPSCs measured in L2/3 PNs were within the range of monosynaptic connections (**Figure 6C**), IPSCs were significantly delayed compared to the EPSCs in the same cells indicating disynaptic inhibition (**Figure 6D**). Indeed, the IPSCs disappeared while the EPSCs persisted after washing in TTX and 4-AP (**Figure 6E**). These observations strongly suggest that excitatory inputs from SSp-bfd drive disynaptic local inhibition onto L2/3 PNs in VISp.

Given the importance of the precise excitation and inhibition balance in sensory perception(Kirkwood, 2015) and its circuit-specific variation, we evaluated the cross-modally evoked E/I balance in L2/3 PNs in VISp. We found that the synaptic input to L2/3 PNs was dominated by the delayed inhibition under different simulation frequencies and durations, which was reflected in the cell-by cell E/I ratio (**Figures 6F, 6G**). Taken together, SSp-bfd directly functionally targets and is able to inhibit L2/3 PNs in VISp.

### SSp-bfd mediates feedforward inhibition via local fast spiking inhibitory neurons in layer 2/3 in VISp

To determine the source of the inhibition evoked via SSp-bfd activation, we compared the functional connectivity between SSp-bfd and L2/3 GABAergic as well as excitatory L2/3 neurons in VISp. For this, we injected AAV.CaMKIIa-hChR2-EYFP in SSp-bfd in GAD/tdtomato transgenic mice to specifically measure cross-modal input to PNs and Interneurons (INs, **Figure 6H**). Moreover, we followed a similar injection approach in PV/tdtomato transgenic mice to gain further subtype input specificity to INs (**Figure 6H**). First, we wondered whether the input from SSp-bfd to different cell types can lead to action potential firing. We measured light-evoked PSPs of neighbouring PNs and INs and observed that both cell types displayed cross-modal input (**Figure 6I**). Strikingly, a fraction of INs fired light-evoked action potentials in response to optogenetic stimulation of SSp-bfd axons. When classifying INs based on their maximum firing frequency obtained by step-current injections (**Figures S5E, S5F**), we found that only fast-spiking interneurons (FS INs) were able to fire action potentials upon blue light stimulation (both in GAD or PV/tdtomato mouse lines, **Figure 6J**). More specifically, while about 50% of FS INs did fire action potentials upon light activation, none of the measured PNs and non-fast spiking INs (nFS INs) showed light-evoked action potentials (**Figure 6K**). These observations strongly suggest that local inhibitory circuitry can be driven by long-range photoactivation and that the strong inhibition observed in VISp is mediated via FS INs indicating they are the main source for the observed feedforward inhibition recruited during cross-modal activation.

We next sought to unravel the reason for FS INs showing light-evoked action potential firing in contrast to PNs albeit using the same power and duration of blue light. In principle, the specific SSp-bfd input strength to FS INs and PNs could differ and consequently explain the ability for one cell type firing action potentials over the other (**Figure 6L**). Alternatively, the biophysical properties of FS INs and PNs targeted by SSp-bfd could vary, resulting in different intrinsic excitability.

To test the first hypothesis, we recorded from neighbouring PNs and FS INs (using the PV/tdTomato mouse line) and measured their light-evoked input strengths (**Figure 6L**). We found that there was no significant difference between the light-evoked EPSC amplitudes in these two cell types (**Figure 6M**). To test the second hypothesis, we measured the intrinsic cell excitability by extracting sub- and suprathreshold electrical properties of PNs and FS INs (both in PV and GAD/tdTomato mouse lines) directly targeted by SSp-bfd. While we found that most subthreshold properties were not significantly different between PNs and FS INs (see e.g. threshold current, **Figure 6N**), most suprathreshold properties were significantly different between these two cell types. Importantly, the difference between the resting membrane and action potential threshold was significantly reduced in FS INs compared to PNs rendering these interneurons more intrinsically excitable (**Figure 6O**). Moreover, the maximal increase of action potential firing frequency between two subsequent current injection steps – a measure of firing gain - was significantly greater for FS INs compared to PNs (**Figures S5G, S5H**). These distinct electrical properties were only found in FS INs but not for nFS INs (**Figure S5H**). In summary, the intrinsic properties of FS INs tend to make these cells more excitable to synaptic input from SSp-bfd compared to PNs.

### Network model identifies a gain-dependent ISN regime to mediate the cross-modal suppression

To investigate whether our optophysiological data can explain whisker stimulation mediated VISp suppression, we built a mathematical cross-modal network model of mean firing-activities of excitatory pyramidal (PN) and fast-spiking inhibitory (FS) neurons in SSp-bfd and VISp (**Figure 7A**). We modeled each cortical area using a well-established canonical cortical recurrent neural network (RNN) (Tsodyks et al., 1998; Wilson and Cowan, 1972) model and connected and constrained them based on our experimental data.

**Figure 7:**
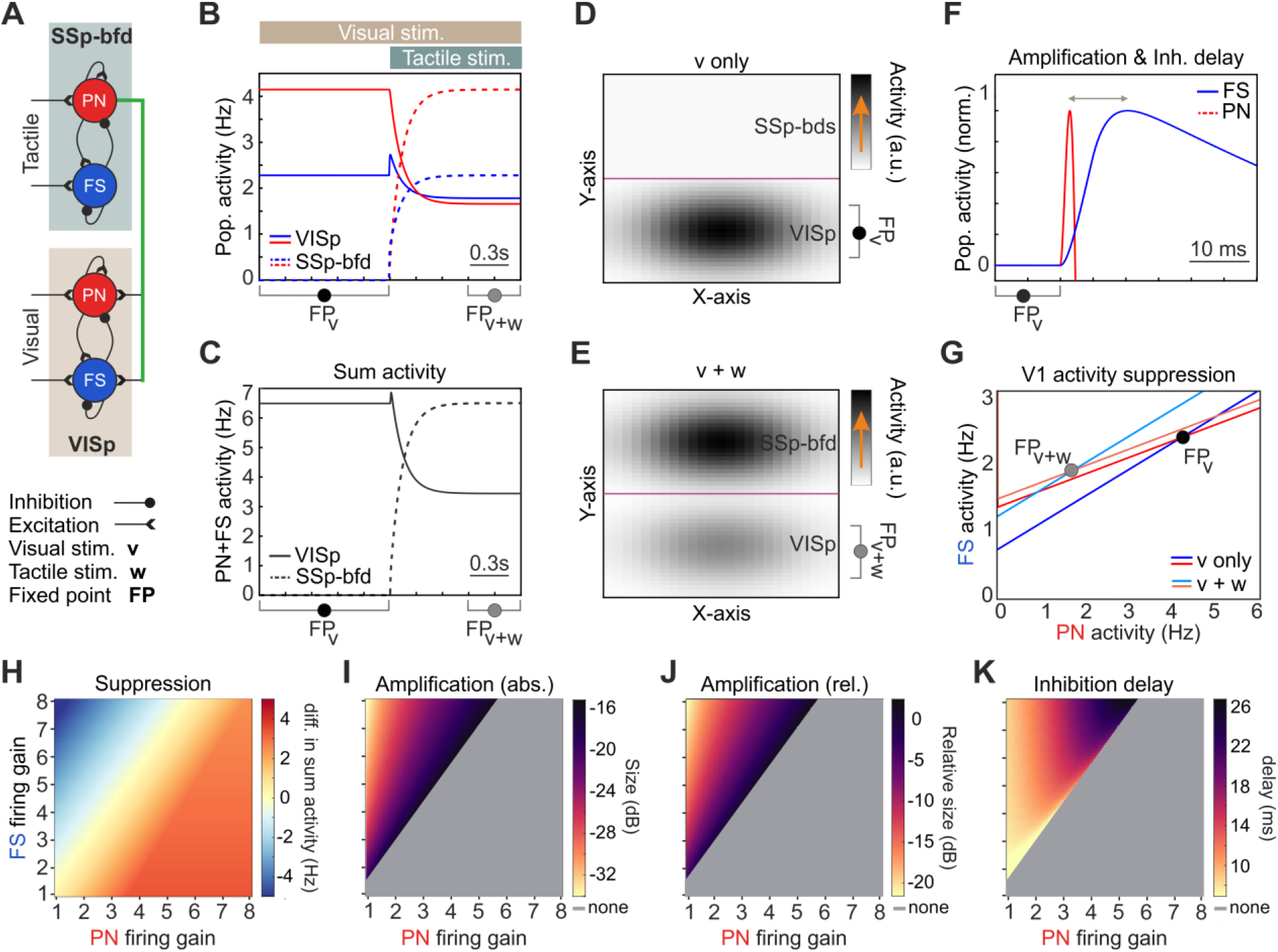
A cortical network model identifies a gain-dependent ISN regime to mediate VISp suppression by tactile stimulation. (A) Schematic diagram of the cross-modal model of coupled VISp and SSp-bfd regions, each modeled as a recurrent neural network (RNN) of pyramidal (PN) and fast-spiking (FS) neural populations. (B) Stimulated visual activity is suppressed by tactile stimulation. Tactile stimulation (w) is added when visual stimulation (v) was already present. These inputs were modeled as constant excitatory inputs. FP_v_ and FP_v+w_ denote the fixed point of VISp-RNN at which its PN and FS population activities (A_PN_ and A_FS_) approached their steady states. (C) Same as (B), but for the sum activities: A_PN_+A_FS_. (D, E) Graphical, spatial illustration of the suppression based on the sum activity levels at FP_v_ and FP_v+w_ shown in (C). (F) SSp-bfd-mediated input to VISp-RNN is transiently amplified by its PN population, as FS response evolve slower (inhibition delay). Zoom-in of the scaled, overlaid VISp-RNN population activities shown in b, around the onset of tactile stimulation. (G) VISp operates, and mediates the cross-modal suppression, under an ISN regime. The A_FS_-A_PN_-plane of the VISp-RNN’s stationary dynamics. The lines represent A_PN_- and A_FS_-nullclines of VISp-RNN’s PN and FS population activities before (v only; red and blue lines) and after adding the tactile stimulation (v+w; brown and bluish lines). Note the reduction in PN and FS activities at FP_v+w_, as compared to FP_v_. (H) The gain-dependency of the strength of VISp suppression by the SSp-bfd-mediated input. Color-coded matrices of the suppression strength computed as the difference in sum activity (FP_v+w_ minus FP_v_), for different combinations of PN and FS firing gains. Negative values (mainly blue) indicate suppression. (I) Same as (H), but for the absolute amplification size, defined in (F). (J), Same as (H), but for the relative amplification size. (K) Same as (H), but for the inhibition delay, defined in (F).

As depicted in **Figures 7B-7C** our model emulates SSp-bfd mediated suppression of visually driven VISp activity. In detail, we first stimulated VISp-RNN and subsequently stimulated the SSp-bfd-RNN using constant excitatory inputs, emulating visual and tactile stimulations. Inputs were sufficiently long allowing the firing activities to reach their steady-states; we call this specific state of each RNN as its fixed point (FP). Strikingly, while the tactile stimulation increases the population activities in SSp-bfd-RNN, it effectively suppresses both PN and FS population activities in VISp-RNN (compare activity levels at FP_v_ and FP_v_+w; **Figure 7B**, solid lines). This suppression is also present in the time course as well as in the spatial illustrations of the steady states of sum activities in these regions (**Figures 7C-7E**). Importantly, these results corroborate our experimental data (see Figure 2 and S2). Moreover, our model shows that SSp-bfd input to VISp is able to be transiently amplified by its PN population (**Figure 7F**), prior to the overall suppression (**Figure 7B**). We further find that this amplification time-window is mainly provided by the slower evolution of FS activity response than PN population, upon the onset of tactile stimulation (**Figure 7F**).

The model suggests that VISp processes the cross-modal input under an inhibition-stabilized network (ISN) regime (Ozeki et al., 2009; Rahmati et al., 2017; Tsodyks et al., 1997), underpinning the suppression phenomenon. This can be seen in A_FS_-A_PN_-plane of VISp-RNN (**Figure 7G**), representing the nullclines (i.e. the steady state dynamics) of its population activities (A_FS_- and A_PN_-nullclines): At the nullclines intersection approached after visual stimulation (FP_v_), both A_PN_- and A_FS_-nullclines have positive slopes but A_FS_-nullcline is steeper which means that inhibition is necessarily and sufficiently strong to stabilize the whole RNN activity (ISN regime). Under this regime both PN and FS populations show an overall reduction in activity after adding tactile stimulation. Note that the suppression under ISN often requires stronger excitatory input to FS, while our optophysiological data show SSp-bfd inputs of similar size to both FS and PN populations in VISp. Under this constraint, our model reveals that the higher firing gain of FS inhibitory neurons compared to PN plays an important role in driving the suppression phenomenon. More specifically, we find that not only the emergence and strength of the suppression (**Figure 7H**), but also the size of transient amplification (**Figures 7I, 7J**) and the inhibition delay (**Figure 7K**) are effectively dependent on this biological constraint. Taken together, our modeling results indicate that VISp processes the SSp-bfd-mediated cross-modal suppression under a gain-dependent ISN regime.

### Tacto-visual convergence in visual proximity space

So far we showed that whisker mediated SSp-bfd activation predominantly suppresses visually driven activity in the anterior part of VISp. This location of postsynaptic neurons is associated with the lower nasal visual space, thereby defining a restricted external space of tacto-visual modulatory influences mediated by SSp-bfd. Here, we investigated the dynamic association of the whisker tips with this modulated visual zone in the mouse proximity space. As demonstrated above, towards their final protraction position, whisker tips also accumulated in the lower, nasal visual space (**Figure 8A**) and this spatial location strongly resembles the visual space coverage of the cross-modally modulated postsynaptic neurons. Indeed, whisker tip locations and the modulated visual space gradually converged during the course of whisker protraction (**Figures 8A, 8B**). Notably, under protraction conditions both, whisker and cross-modally modulated visual space practically covered the same spatial extent and were strongly overlapping (**Figure 8A**). Moreover, in this constellation practically all caudal whiskers identified to be important for tactile integration in VISp (see Figure 4) were located inside the modulated visual space (**Figures 8A, 8B**). This suggests that mice can actively move their whiskers through the space in which visual processing is modulated by the whiskers themselves. This further reveals that tacto-visual convergence in mouse proximity space is precisely reflected at the cortical level of VISp.

**Figure 8:**
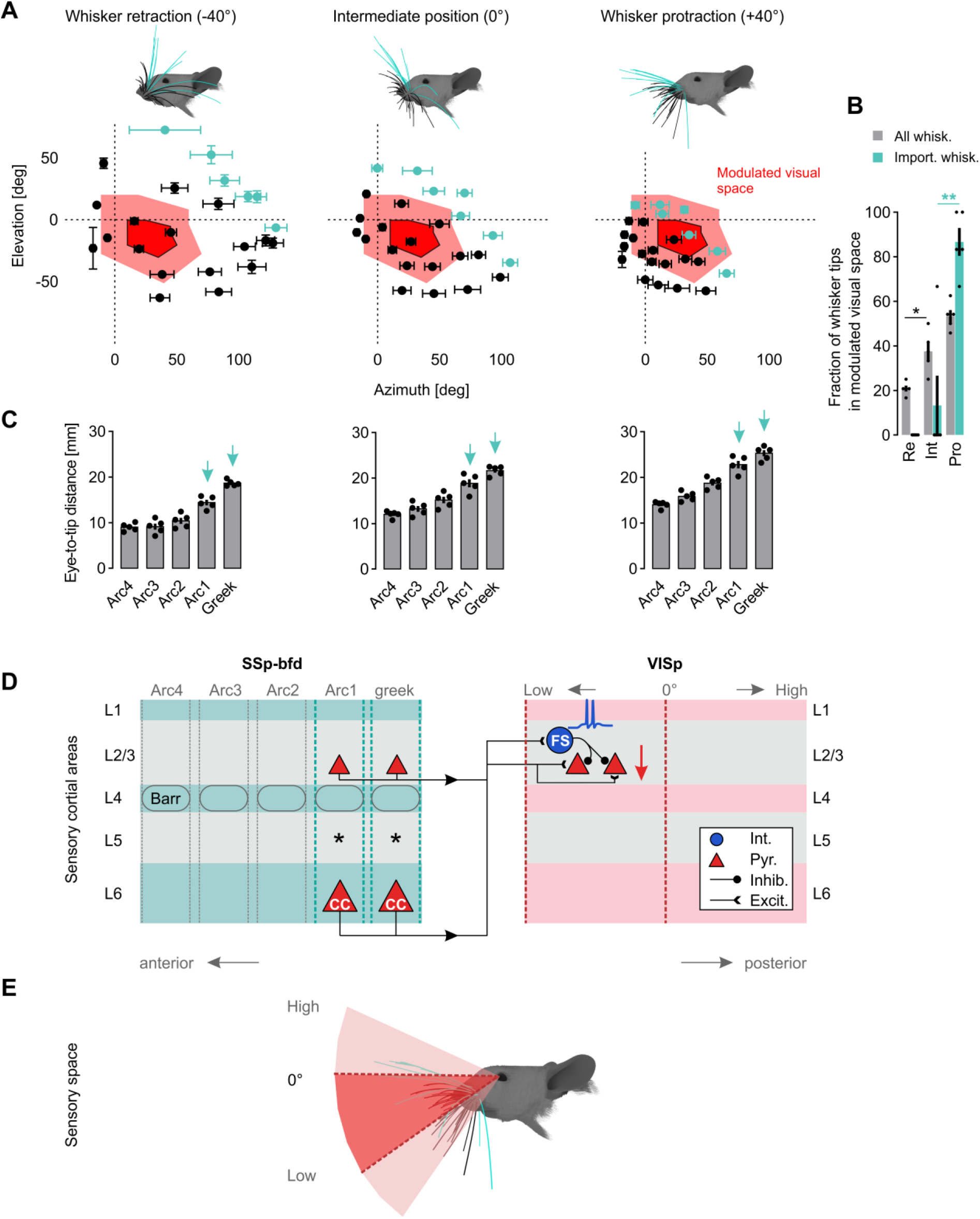
Tacto-visual convergence in the mouse proximity space. (A) Mapping of tacto-visual overlap along azimuth and elevation. Red area with surrounding solid black line: Coverage map of visual space covered by the VISp region with the highest fraction (at least 5% on average) of postsynaptic neurons labeled after SSp-bfd injections. Bright red area: Coverage map of visual space covered by the same VISp region mapped by eye movements. Centroids indicate positions of whisker tips. Colored centroids represent the tips of the caudal whiskers above identified to be important for visual processing. (B) Fraction of whisker tips located within the visual space covered by VISp under eye movement conditions (bright red area in (A)) during whisker retraction (Re), in the intermediate position (Int) and during protraction (Pro). Black circles indicate data points of individual mice and bars indicates the mean fraction of whisker tips on total number whiskers ± s.e.m (the 24 large whiskers, grey bars, Re vs. Int, p=0.0331; Int vs. Pro, p=0.1349; the 7 “important” whiskers, colored bars, Re vs. Int, p=1; Int vs. Pro, p=0.0087; paired t-tests followed by Bonferroni correction). (C) Diagrams show the average eye-to-whisker tip distance of each of the arcs under retraction, intermediate and protraction conditions ± s.e.m. Black dots represent data points of individual mice. Colored arrows indicate the arcs containing the whiskers important for cross-modal tactile integration in VISp. *p<0.05, **p<0.01. (D) Summary Figure. In SSp-bfd both excitatory CC-neurons in L6 and excitatory neurons in L2/3 predominantly located in posterior barrel columns of SSp-bfd send direct axonal projections to the anterior region of VISp. Here, these projections innervate excitatory pyramidal and FS inhibitory neurons in L2/3. Because this innervation evokes spiking in FS but not pyramidal neurons, visually evoked neuronal activity in pyramidal neurons becomes inhibited (feed-forward inhibition). (E) Summary Figure. The anterior region of VISp processes visual signals originating in the lower, nasal visual space. Under whisker protraction conditions such as during object exploration, whisker tips are precisely located in this part of the visual space. Thus, tacto-visual convergence in VISp is precisely reflected in the mouse proximity space. Colored whiskers predominantly innervate the posterior barrel columns and thus play an important role in SSp-bfd mediated suppression of VISp activity. Their tips reach furthest into the mouse proximity space increasing the probability that they are involved in the first contact with an object during exploratory behavior.

However, what might be a reason for the higher relative importance of the caudal whiskers for tactile integration in VISp? Measuring their length and eye-to-tip distance in our 3D model revealed that these whiskers are the longest, whose tips always reach furthest away from the eye into the proximity space (**Figure 8C**). Thus, when mice approach objects there is a high probability that these particular whiskers are involved in the first contact. Thereby, visual signals originating from the same object may be suppressed instantly.

## 3 Discussion

Here we find that whisker stimulation suppresses visually elicited responses in VISp via a SSp-bfd originating cortico-cortical pathway that drives FS inhibitory neuron mediated feed-forward inhibition (FFI) in L2/3 excitatory neurons (**Figure 8D**). We show that both projection and postsynaptic neurons involved in this microcircuit are predominantly located at the border regions of SSp-bfd and VISp, respectively, which are in close proximity to each other. Thus, in terms of visual field representations of postsynaptic neurons in VISp, SSp-bfd mediated suppression is likely to be restricted to the lower, nasal part of the visual space. Importantly, this space substantially overlaps with the external space, where whiskers perform tactile exploration (**Figure 8E**). This suggests that the specific tacto-visual convergence in proximity space is also reflected at the anatomical and functional level of VISp, providing a cortical anatomical locus for sensory interactions.

Multisensory convergence has been suggested to mainly occur in higher-order cortical association areas (Felleman and Van Essen, 1991; Jones and Powell, 1970; Sugihara et al., 2006; Teichert and Bolz, 2018). However, evidence shows that even primary cortical sensory areas, in which sensory processing was previously assumed to be conducted on a sense-by-sense basis (Wallace et al., 2004), can be multisensory (Deneux et al., 2019; Ibrahim et al., 2016; Iurilli et al., 2012; Kayser et al., 2008; Renard et al., 2022; Teichert and Bolz, 2017, 2018; Velez-Fort et al., 2018). For example, visual processing in mouse VISp is influenced by sounds (Deneux et al., 2019; Ibrahim et al., 2016; Iurilli et al., 2012; Teichert and Bolz, 2017; Teichert et al., 2017). While tacto-visual convergence in rodents has been suggested to occur in the superior colliculus (Drager and Hubel, 1975; Meredith and Stein, 1983; Shang et al., 2019) and higher-order cortical areas (Nikbakht et al., 2018; Olcese et al., 2013), it remained unclear whether primary sensory areas contribute to tacto-visual processing as well. Here, we demonstrate that visually evoked responses in VISp are also cross-modally modulated by the sense of touch. In line with our findings, cross-modal suppression is a frequently observed feature of multimodal tactile integration in sensory cortical areas both in humans (Ide et al., 2016) and other species (Meredith and Allman, 2015; Rao et al., 2014). For instance, whisker stimulation causes a global suppression of sound evoked activity in mouse AUDp (Lohse et al., 2021). Thus, tactile stimulation induced cross-modal suppression of sensory elicited activity in other cortical primary sensory areas may represent a mechanism by which tactile inputs from nearby objects that require immediate attention are processed with priority. This interpretation is in line with our finding that especially the long caudal whiskers covering the largest search space during whisking are particularly important for VISp processing (**Figures 4** and **8**). These whiskers are likely involved in the first contact with an object especially during navigation and object exploration, when whiskers are protracted (Sofroniew and Svoboda, 2015). This will lead to immediate VISp suppression and may therefore shift attention towards tactile cues. However, although tactile driven suppression of visual activity in VISp may reflect the higher relative importance of tactile cues in close proximity, a reduction of VISp responses could be accompanied by a suppression of non-specific noise (Nikbakht et al., 2018) in the visual stimulus representation potentially sharpening visual tuning. Indeed, auditory stimulation sharpens visual orientation selectivity in VISp (Ibrahim et al., 2016). Taken together, our data strongly support the growing concept of multisensory primary sensory cortical areas. Thus, along sensory information streams, these early hierarchical areas potentially belong to the first brain regions where multisensory integration occurs suggesting that downstream higher-order cortical areas and association areas receive a preprocessed digest of multisensory information.

It has been described that optimal multisensory integration is achieved when sensory stimuli are presented to different sensory modalities in not only a temporally but also spatially coherent manner (Holmes and Spence, 2005). In terms of the latter “spatial rule” (Holmes and Spence, 2005) we find that the anatomic arrangement of whiskers and eyes ensures that objects in the mouse proximity space are usually sensed through both of these modalities. According to our simulations, this is especially true when mice protract their whiskers (**Figures 1** and **8**). In this condition, an object in the lower visual field on the ground might be palpated more efficiently by more whiskers leading to a strong activation of the (posterior) barrel columns in SSp-bfd and ultimately to a stronger cross-modal innervation of VISp. In conclusion, the marked peripheral overlap of tactile and visual information streams may allow for optimal tacto-visual integration in VISp.

It has been suggested that primary sensory areas are separated from one another by transitional multisensory zones as revealed by *in vivo* electrophysiological recordings in rats (Wallace et al., 2004). Indeed, in rodents, the higher-order visual area RL, a part of the rodent posterior parietal cortex (Hovde et al., 2019; Nikbakht et al., 2018), is situated between SSp-bfd and VISp and represents a transitional multisensory area converging both tactile and visual signals (Nikbakht et al., 2018; Olcese et al., 2013). Our results extend this view by demonstrating that such transitional zones even exist within primary sensory areas. We find that projection neurons in SSp-bfd and postsynaptic neurons as well as SSp-bfd originating axons in VISp display strong gradients pointing away from each other (**Figures 3** and **5**). This suggests that SSp-bfd exerts its suppressive influence mainly on neurons located in the anterior part of VISp. Consequently, the vicinity of projection and postsynaptic neurons may ensure a fast and economic integration of tactile signals in VISp. Taken together, these findings indicate that mainly transitional border regions of primary sensory areas integrate cross-modal sensory information from neighboring sensory cortical regions. These results might have further implications for defining subdivisions even within primary sensory cortical areas, to distinguish between uni- and multisensory subregions, based on anatomy and function.

Interestingly, neurons in the anterior transitional border region of VISp display distinct functional features as compared to neurons located more posterior. In mice these neurons are significantly activated by objects in close proximity (near disparity tuned) (La Chioma et al., 2019). Given the strong overlap of the visual space with the space mapped by the whiskers (**Figures 1** and **8**), these objects may also be in reach of the whiskers. Moreover, neurons representing the lower visual field are significantly more responsive to coherent visual motion (Sit and Goard, 2020). Interestingly, also whiskers and their corresponding neuronal representations in SSp-bfd act as motion detectors (Laboy-Juarez et al., 2019). Thus, the integration of tactile signals in visual processing in the anterior region of VISp may create a multisensory representation of both peripersonal space and moving objects within this space. This is potentially important for multisensory behaviors requiring tacto-visual interactions such as object exploration (Nikbakht et al., 2018) or predatory hunting (Shang et al., 2019).

Our imaging data indicate a more global suppression of visually driven VISp responses beyond the anterior part of VISp (**Figure 2**). This might be explained by the limited spatial and temporal resolution of intrinsic signal imaging, or by a slow and delayed horizontal spread (Das and Gilbert, 1995; Fehervari et al., 2015; Orbach and Van Essen, 1993) of the multisensory signals from the anterior to posterior part of VISp. Alternatively, as RL and some other higher-order visual areas contain numerous SSp-bfd-recipient postsynaptic neurons (**Figure 5**) and send strong feedback projections to VISp (Glickfeld and Olsen, 2017; Wertz et al., 2015), we cannot rule out that these areas provide additional whisker information to posterior VISp.

Audiovisual processing in VISp is mediated by direct cortico-cortical connections from AUDp to VISp (Deneux et al., 2019; Ibrahim et al., 2016; Iurilli et al., 2012). For example, optogenetic stimulation of AUDp projections modulates visually evoked responses in VISp (Iurilli et al., 2012). Similarly, our anatomical and functional data strongly argue that also tacto-visual integration is achieved by the recruitment of direct cortico-cortical connections originating in SSp-bfd (**Figure 3**). On the other hand, our data do not provide evidence for a pathway in which SSp-bfd relays whisker information to VISp-projecting subcortical areas (**Figure 5**), as recently suggested for tactile integration in AUDp (Lohse et al., 2021). These results support the novel view of anatomically and functionally interconnected primary sensory cortical areas. Strikingly, we observe that VISp projecting neurons in SSp-bfd are most abundant in L6 (**Figure 3**). This is in line with previous retrograde tracing studies in rats (Bieler et al., 2017). In contrast, by using the b-fragment of cholera toxin in mice (Masse et al., 2017) for retrograde tracing experiments, previous investigations revealed that projection neurons involved in the same pathway are mainly located in L5 and only a small portion was found in L6 (Masse et al., 2017). This discrepancy might be explained by the limited depth of cortical injections performed by these authors (Masse et al., 2017). In addition, the usage of newly developed highly efficient tracers such as retro-AAVs (Tervo et al., 2016) as used here, reveals a more prominent involvement of L6 in cortico-cortical communication (Liang et al., 2021), than previously thought.

Notably, L6 is located in a strategic position within the cortex, receiving input from and providing output to the local column (Velez-Fort et al., 2014) and other cortical and sub-cortical brain regions (Briggs, 2010; Liang et al., 2021). However, little is known about the function and projection patterns of CC neurons in L6. L6CCs in sensory areas receive strong thalamic input (Crandall et al., 2017) and send extensive and horizontally orientated projections (Velez-Fort et al., 2014) to cortical motor areas (Zhang and Deschenes, 1997) and across the corpus callosum (Liang et al., 2021). Here, we extend this knowledge by demonstrating that L6CC projections can also cross the border of their host primary sensory area to innervate primary areas of other sensory modalities. This suggests that L6CCs are key players in cross-modal integration. Importantly, neuronal responses in L6CCs in rodent SSp-bfd to whisker deflections precede those in all other excitatory cell types by 3 ms (including neurons in L4) (Egger et al., 2020). Thus, these neurons are ideal candidates to rapidly and efficiently relay whisker information to VISp for tacto-visual integration as they are operating on short latency time scales.

Our optophysiological data in combination with our mathematical cortical network model show that cross-modal SSp-bfd mediated suppression of VISp activity can be largely explained by recruitment of FFI onto L2/3 PNs (**Figures 6** and **7**). This observed long-range recruitment of L2/3 for multisensory integration is in line with numerous previous studies (Deneux et al., 2019; Ibrahim et al., 2016; Iurilli et al., 2012). Long-range connections from different brain areas have been shown to preferentially recruit specific sets of L2/3 INs in a given postsynaptic brain area (Naskar et al., 2021). We find that FFI is mediated by local PV^+^ FS cells, which have been shown to be the most abundant neuron type in FFI (Tremblay et al., 2016). The previously described perisomatic targeting of PV^+^ INs together with the here observed intrinsic properties enabling high speed fidelity, provides unique temporal filtering properties permitting precise coincidence detection onto postsynaptic PNs. However, other layers and interneuron types have also been shown to be involved in FFI and ultimately in multisensory integration (Ibrahim et al., 2016; Iurilli et al., 2012). Therefore, the exact circuit motif for long-range cross-modal FFI might be specific for a given pair of pre- and postsynaptic cortical brain areas. Additionally, the specific circuit recruitment and the gain of involved neurons has been shown to vary based on internal and external influences (Ferguson and Cardin, 2020). Thus, the effects of whisker stimulation on VISp responses may dynamically adapt (**Figure 7**) or even inverse (e.g. excitation) in response to changing inputs depending on influences such as arousal, attention, locomotion or specific stimulus features (Ferguson and Cardin, 2020).

In summary, our study provides direct anatomical and physiological evidence for multisensory integration at the level of primary sensory cortices where external space is shared by two sensory systems. It further suggests that primary sensory cortices are heavily involved in generating complex multisensory representations for high-order processing.

## 4 Methods

### Animals

All experiments were performed on 4–14 week old mice of both sexes. The following mouse strains and transgenic mouse lines were used: C57BL/6J and Ai14 (Cre-dependent tdTomato reporter) mice (Jackson Laboratories, RRID: IMSR_JAX:007914). Ntsr1-Cre-tdTomato reporter mice were bred by crossing Ntsr1-Cre (layer 6 cortico-thalamic cells; GenSAT 030648-UCD) with Ai14 mice. Similarly, Gad2-Cre-tdTomato and PV-Cre tdTomato reporter mice were bred by crossing Gad2-IRES-Cre with Ai14 mice. Mice were raised in standard cages on a 12 h light/dark cycle, with food and water available *ad libitum*. Animal housing is regularly supervised by veterinaries from the state of Thuringia, Germany. All experimental procedures were in accordance with the German Law on the Protection of Animals, the corresponding European Communities Council Directive 2010 (2010/63/EU), with the UK Home Office regulations (Animal Scientific Procedures, Act 1986), approved by the Animal Welfare and Ethical Review Body (AWERB; Sainsbury Wellcome Centre for Neural Circuits and Behaviour) and in compliance with ARRIVE guidelines. Every effort was made to minimize the number of animals and their suffering.

### 3D-reconstruction of the mouse whisker array and the visual space

Because the reconstructed whisker array was finally aligned and fit to an existing realistic 3D model of the mouse, which was generated from 8-13 weeks old female mice (Bolanos et al., 2021)(https://osf.io/h3ec5), we used mice of the same age and sex for whisker reconstruction. We always reconstructed the 24 large caudal whiskers (greeks, and arcs 1 to 4) on the left side of the snout. For stability reasons, whiskers were reconstructed from dead mice approximately 2 h after death initiated by an overdose of isoflurane in a sealed container. The 3D distribution of the whiskers in this condition was defined as their intermediate position. Importantly, this position only marginally differed from the position of whiskers in anesthetized mice (data not shown). For fixation the scalp was removed and a small magnet was glued to the skull using cyanoacrylate. This magnet was then attached to a metal bar fixed in a micromanipulator (BACHOFER, Reutlingen) to allow for adjustments of the mouse position. Whiskers were then reconstructed by stereo photogrammetry (Heist et al., 2018; Stark et al., 2019). In brief, the mouse head including the whiskers was illuminated by structured light generated by a custom-written software in MATLAB 2019-2022 and delivered by a commercial projector (AOPEN QF12 LED Projector). Subsequently 90 stereo images were taken by two cameras (ALLIED Vision Technologies, guppy pro) focusing on the mouse head from different angles. This procedure was repeated 4-6 times with the mouse positioned in different orientations to ensure capturing of the whole whisker array. Finally, the detection of corresponding homologous point pairs then allowed for 3D-reconstruction of the mouse head including the whiskers via triangulation using custom software written in C++11. Generated point-clouds were then visualized, aligned and processed using the free open-source software CloudCompare v2.12 (https://www.cloudcompare.org) to obtain one final 3D point cloud including all whiskers of the array. However, occasionally some whiskers were not fully reconstructed up to their thin tip. To solve this issue all whiskers of the same previously scanned mice were cut right above the skin, micrographs were taken using a standard stereo microscope (Stemi SV6, ZEISS) and whiskers were traced in 2D along their midline from their base to their tips using the “b-spline” tool in CorelDRAW 2021.5. Notably, after cutting the whiskers additional stereo images (90) were taken from the mystacial pad after marking the whisker base points with ink for better visualization and a 3D-reconstruction was performed as described above. Thus, the exact origin of each individual whisker could be determined. Traced whiskers were then scaled to their real size and imported into CloudCompare v.2.12 as .obj-files. Here, corresponding 3D-reconstucted and 2D traced whiskers were first manually aligned and then finely registered to each other using the iterative closest point (ICP)-algorithm included in CloudCompare v.2.12. The alignment of the 3D-reconstructed and 2D-traced whiskers was considered sufficient as whisker curvature has been observed to occur mostly in one plane (Dougill et al., 2020). Indeed, the average radial distance between these two representations ranged only around 0.16±0.1 mm across whiskers of all mice. Finally, the head of the existing realistic 3D model of the mouse (Bolanos et al., 2021) was imported from the free open-source software Blender v3.2.0 (https://www.blender.org/download/) into CloudCompare v.2.12 as an .obj-file and the 3D-reconstructed point clouds of the mouse head including the traced whiskers and the mystacial pad were aligned to it, based on visual inspection. Thus, whiskers now had the correct origin, orientation, length and shape with respect to the realistic model of the mouse head (Bolanos et al., 2021). Subsequently, this model of the whisker array was imported back into Blender v3.2.0 as a .dxf-file and re-adjusted to the realistic 3D mouse model.

The 3D reconstruction of the visual space coverage of VISp was performed based on retinotopic mapping data from (Zhuang et al., 2017). These maps contain the azimuth and elevation as mapped across the visual cortical area by presenting spherically-corrected checkerboard visual stimuli drifting across the visual field (Zhuang et al., 2017). Maps of mouse VISp containing azimuth and elevation contour plots (contours of 5° intervals from 0° to 90° in azimuth and −25° in elevation) (Zhuang et al., 2017) were then used to estimate the extend of VISp visual space coverage. For this, azimuth and elevation coordinates along the border of VISp were determined and used for 3D-reconstruction of visual space in Blender v3.2.0. Here, we created a left eye-centered spherical coordinate system and implemented the azimuth and elevation values of the VISp visual space coverage. The resulting visual space was cut at 3 cm away from the left eye for better illustration.

For quantifying the overlap between EGFP-expressing projection neurons and GAD-tdTomato or Ntsr1-tdTomato expressing cells in SSp-bfd, 50 μm-thick brain slices were mounted in Vector Shield mounting solution. Coronal images across slices were then acquired on a confocal microscope (Leica SP8, 10x oil immersion, optical section step of 1 μm) and overlap was manually quantified in ImageJ.

### Simulation of whisker and eye movements

Simulations of whisker retraction and protraction were performed in Blender v3.2.0 (https://www.blender.org/download/). Generally, in mice, during retraction and protraction the azimuthal whisker angel is highly correlated across whiskers and elevation movements are anticorrelated with azimuth (Petersen et al., 2020). In other words, when mice protract their whiskers they simultaneously move them downwards and when they retract them they move upwards with respect to the mouse head. From these data we estimated, that downwards and upwards movements of all whiskers roughly follow a plane fitted through the basepoints (on the mystacial pad) and whisker tips of the corresponding row (but see (Petersen et al., 2020), Fig. 4E). Such a plane was created for the whiskers of each row in blender v3.2.0 using a custom-based software written in Phyton (https://raw.githubusercontent.com/varkenvarken/blenderaddons/master/planefit.py). In order to simulate extreme whisker retraction and protraction, whiskers were rotated around their basepoint parallel to this plane to −40° or +40° as starting from their intermediate position. The most caudal creek whiskers (α-δ) were rotated parallel to the plane of their corresponding rows (A-D). Typically, during retraction and protraction whiskers also rotate around their longitudinal axis (Petersen et al., 2020). However, as rotation would only marginally affect the position of the whisker tips in space, this movement was not included in our model. Animations of simulated whisking behavior were created in Blender v.3.2.0 and rendered using the Eevee-engine. Eye-movements were simulated by extending the VISp visual field coverage by 20° in each direction.

### Intrinsic signal imaging

Animals were initially anesthetized with 4% isoflurane in a mixture of 1:1 O2/N2O and placed on a heating blanket (37.5 °C) for maintaining body temperature. Subsequently, mice received injections of chlorprothixene (20 μg/mouse i. m.) and carprofen (5 mg/kg, s. c.). The inhalation anesthesia was applied through a plastic mask and maintained at 0.5 - 0.7% during experiment. The skin above the right hemisphere was removed and a metal bar was glued to the skull to fix the animal in a stereotaxic frame using dental acrylic. Next, the skin above the left hemisphere was removed to expose visual and somatosensory cortical areas. The exposed area was covered with 2.5% agarose in saline and sealed with a glass coverslip. Cortical responses were always recorded through the intact skull.

Using a Dalsa 1M30 CCD camera (Dalsa, Waterloo, Canada) with a 135 × 50 mm tandem lens (Nikon, Inc. Melville, NY), we first recorded images of the surface vascular pattern via illumination with green light (550 ± 2 nm) and, after focusing 600 μm below the pial surface, intrinsic signals were obtained via illumination with red light (610 ± 2 nm). Frames were acquired at a rate of 30 Hz and temporally averaged to 7.5 Hz. The 1024 × 1024 pixel images were spatially averaged to a 512 × 512 resolution. Responses of VISp were recorded as described originally (Kalatsky and Stryker, 2003). Briefly, the method uses a periodic stimulus that is presented to the animal for some time and cortical responses are extracted by Fourier analysis. In our case, the visual stimulus was a drifting horizontal light bar of 2° width, a spatial frequency of 0.0125 cycles/degree, 100% contrast and a temporal frequency of 0.125 Hz. It was presented on a high refresh rate monitor (Hitachi Accuvue HM 4921-D) placed 25 cm in front of the animal. Visual stimulation was adjusted so that it only appeared in the nasal visual field of the left eye (−5° to +15° azimuth, −17° to +60° elevation). The stimulus was presented to both eyes simultaneously for 5 min. Thus, it was repeated for about 35 times during one presentation. The facial whiskers on the left side of the snout were first stimulated by a moving metal pole which was connected to an Arduino (Arduino-Uno, Genuino, USA) controlled hybrid polar stepping motor (SOYO, USA). Simultaneously, the metal pole vibrated with a frequency of 20 Hz (sinusoidal). The pole was first moved in dorsoventral direction from the A-row to the E-row of the whiskers array within 8 s (temporal frequency of 0.125 Hz) for 5 min, thereby sweeping over the whisker tips and deflecting them by an angle of about 20-25° before they whipped back in their normal position. In order to remove the hemodynamic delay of the intrinsic signals, both the visual and whisker stimulus was reversed in the following presentation period.

For simultaneous imaging in both SSp-bfd and VISp, we synchronized the visual and whisker stimulus temporally and spatially. In detail, as the light bar started moving from the bottom of the monitor (−15°), the metal bar started at the same time at the A-row of the whisker array. During the following 8 s the light bar moved to the top of the monitor meanwhile the metal bar moved in dorso-ventral direction towards the E-row of the whiskers. The synchronization was also maintained after the stimulus reversal. To investigate whether this bimodal sensory stimulation affects VISp or SSp-bfd activity, we performed imaging under unimodal visual stimulation in the same mice. Uni- and bimodal stimuli were presented in pseudorandom manner. Experiments in that we only stimulated the whiskers were performed in complete darkness.

From the recorded frames the signal was extracted by Fourier analysis at the stimulation frequency and converted into amplitude and phase maps using custom software (Kalatsky and Stryker, 2003). For data analysis we used MATLAB 2019-2022. In detail, from a pair of the upward and downward maps (visual or somatosensory), a map with absolute visuotopy or somatotopy and an average magnitude map was computed. The magnitude component represents the activation intensity of VISp or SSp-bfd. Since high levels of neuronal activity decrease oxygen levels supplied by hemoglobin and since deoxyhemoglobin absorbs more red light (610±2nm), the reflected light intensity decreases in active cortical regions. Because the reflectance changes are very small (less than 0.1%), all amplitudes are multiplied with 10^4^, so that they can be presented as small positive numbers. Thus, the obtained values are dimensionless. For each stimulation condition we recorded at least three magnitudes of VISp (or SSp-bfd) responsiveness and averaged them for data presentation.

In another set of experiments whisker stimulation was performed using air puffs generated by a picospritzer. The air puffs were applied with a frequency of 2 Hz and a duration of 500 ms through a hollow needle directed to the whiskers on the left side of the snout from frontal. Great care was taken to direct the air puffs only to the whiskers and to avoid any stimulation of the fur on the mouse’s head and body. Air puffs induced whisker deflections with an angle of about 20-25°. We recorded VISp responses in the absence and presence of air puffs. Cortical responses were again extracted by Fourier analysis as described above.

To examine whether the moving metal pole *per se* causes visual disturbances, we trimmed the whiskers on the left side of the snout in another group of mice using fine scissors and recoded VISp activation induced by the moving light bar in the absence and presence of the moving metal pole. Cortical responses of VISp were extracted by Fourier analysis as described above.

In another group of mice, we investigated whether the sound created by the air puffs cross-modally affected visually evoked VISp responses. For this, we again trimmed whiskers on the left side of the snout. Thus, applying air puffs (from the same frontal position as above) did not lead to whisker stimulation. We recorded VISp responses in the absence and presence of air puffs. Cortical responses were extracted by Fourier analysis as described above.

In another subset of experiments we investigated whether the loss of afferent whisker input to the brain cross-modally affects VISp responses. For this, we cut the infraorbital nerve (ION) on the left side of the snout by gently opening the skin using fine forceps and cutting the ION using fine scissors. A sham surgery was performed by opening the skin above the ION. Then the nerve was gently touched by the fine scissor but left intact. VISp responses were recorded before and after the sham surgery and the ION cut under simultaneous whisker stimulation using air puffs as described above.

### Immunohistochemistry

The number of c-fos positive cells in different layers of VISp was examined in control mice with normal whiskers and mice with bilaterally removed whiskers (WD). Firstly, awake animals were dark adapted for 2 h. Subsequently, in WD mice all macrovibrissae were trimmed using an electric shave. This typically took 1-2 min. In control mice whiskers were sham trimmed by moving the electric shave through the whiskers while switched off. After this, mice were placed in and enriched environment for bimodal visual and somatosensory (in control mice) stimulation for 1.5 h. The environment was surrounded by four monitors showing a moving sine wave grating (0.1 cyc/deg, 100% contrast) for boosting visual stimulation. Simultaneous whisker stimulation (in control mice) was achieved by obstacles (paper roles, wood wool, brushes) placed on the bottom of the environment. Mice remained here for 1.5 h. Subsequently, animals were deeply anesthetized by an intraperitoneal injection of a ketamine (100 mg/kg)/xylazine (16 mg/kg) solution. Animals were perfused transcardially using 100 mM phosphate buffered saline (PBS) followed by 4% paraformaldehyde (PFA) in PBS. The removed brains were post fixed in 4% PFA and cryoprotected in 10% and 30% sucrose in PBS. Frozen sections of 20 μm thickness were taken using a cryostat (Leica).

For 3,3’-diaminobenzidine (DAB) stainings sections were washed in PBS containing 0.2% TritonX and subsequently a peroxidase block (1% H_2_O_2_ in PBS) was performed for 30 min. After blocking in 10% serum, 3% bovine serum albumin (BSA), and 0.2% TritonX-100 in PBS for 1 h, sections were incubated free floating with a primary antibody against c-fos (rabbit-anti-c-fos, 1:250, Santa Cruz) over night and at room temperature. After 1 h of incubation with a biotinylated secondary antibody (goat-anti-rabbit, 1:1000, Vector) at room temperature, sections were washed in PBS. The DAB reaction usually took about 15 min and was performed in the absence of light. The reaction was stopped with PBS, the stained sections were embedded in mowiol (Roth) and sealed with a coverslip. The sections were observed using a bright field microscope (Olympus) using a 10× objective and analyzed with ImageJ. We always used 4–5 sections containing the anterior VISp or SSp-bfd of each animal and averaged the number of stained c-fos positive nuclei within the specific cortical area. The cortical region of the mouse visual and somatosensory cortex was determined on atlas basis (Wang et al., 2020)

For fluorescence immunohistological stainings cryosections were postfixed in 4% PFA for 30 min, washed in 0.2% TritonX-100 in PBS, blocked in 10% serum, 3% BSA, and 0.2% TritonX-100 in PBS for 1 h, followed by the incubation with the primary antibodies overnight. After washing, slices were incubated with the secondary antibody for 2 h. Following primary antibodies were used: rabbit-anti-c-fos (1:200, Santa Cruz), rabbit-anti-PV (1:1000, Abcam; directly labeled to Alexa 488 using a Zenon Rabbit IgG Labeling Kit) and goat-anti-SOM (1:100, Santa cruz). Following secondary antibodies were used: Alexa488 donkey-anti-rabbit (1:1000, Jackson Immuno Research), Cy3 donkey anti goat (1:1000; Jackson Immuno Research), Cy3 goat-anti-rabbit (1:1000, Jackson Immuno Research). After embedding slices in mowiol (Roth), pictures were scanned with a LSM510 (Zeiss) using a 20× objective and analyzed with ImageJ. We used 4–5 sections of the anterior VISp per animal and counted the number of PV or SOM positive interneurons within VISp. This value was averaged throughout the 4–5 sections to obtain one value per animal. Next we counted the numbers of PV and SOM positive cells which also expressed c-fos. Thus, we could calculate the percentage amount of double-stained cells of all PVs or SOMs, respectively. Double labeled cells were always counted within single focal planes along the z-axis. During counting, we switched between the green (PV or c-fos) and red channel (SOM or c-fos) to ensure that we only took cells and nuclei at the same location and with a proper staining into count. Furthermore, we only counted cells which clearly displayed a typical cellular shape and size. The location of VISp was determined based on the Allen Reference Atlas (coronal, 2D) (Wang et al., 2020).

### Stereotaxic viral injections

All surgical procedures, including the craniotomies, and virus injections, were carried out under isoflurane (2%–5%) and after carprofen (5 mg/kg, s.c.) had been administered. For virus injections, mice were anaesthetized under isoflurane (~2%) and craniotomies performed. Virus injection was performed using borosilicate glass injection pipettes (Wiretrol II; Drummond Scientific) pulled to a taper length of ~30 mm and a tip diameter of ~50 μm. Viruses were delivered at a rate of 1–2 nl/s using Nanoject III (Drummond Scientific, USA). Viruses were injected at three cortical depths covering all layers of the VISp and SSp-bfd, respectively. After injections, the craniotomy was sealed with silicon (kwik-cast), the skin was resutured and animals were allowed to recover for 2–4 weeks. Injection coordinates for SSp-bfd and VISp were based on the Allen Reference Atlas (coronal, 2D) (Wang et al., 2020). For retrograde viral tracing we injected rAAV2-retro-EF1a-H2B-EGFP (Nuclear retro-AAV, titer: 8.8 x 10^13^ GC per ml) or rAAV2-retro.CAG.GFP (Cellular retro-AAV, titer: 5 × 10^13^ GC per ml). For anterograde tracing, we injected AAV2/1-hSyn-Cre (titer: 1.3 × 10^13^ GC per ml). For functional connectivity experiments in acute slices, we injected either a mixture of 1:1 of AAV2/1-hSyn-Cre and AAV-2/1-CaMKIIa-hChR2(H134R)-EYFP or a mixture of 1:1 of AAV-2/1-CaMKIIa-hChR2(H134R)-EYFP and saline.

For perfusions, mice were first deeply anaesthetized using a ketamine (200 mg/kg)/xylazine (20 mg/kg) mixture. A blunt needle was placed in the left ventricle, whilst an incision was performed in the right atrium of the heart. Following this, blood was first cleared using 100 mM PBS. Subsequently, the animal was perfused with saline containing 4% PFA. After successful fixation, the head was removed and the brain dissected out. The brain was further fixed in 4% PFA overnight at 4 °C, and then stored in 100 mM PBS at 4 °C until ready for imaging.

### Brain wide serial two-photon imaging

For serial section two-photon imaging, on the day of imaging, brains were removed from the PBS and dried. Brains were then embedded in agarose (4%) using a custom alignment mould to ensure that the brain was perpendicular to the imaging axis. The agarose block containing the brains were trimmed and then mounted onto the serial two photon microscope containing an integrated vibrating microtome and motorized x–y–z stage (Osten and Margrie, 2013; Ragan et al., 2012). For this, a custom system controlled by ScanImage (v5.6, Vidrio Technologies, USA) using BakingTray (https://bakingtray.mouse.vision/) was used. Imaging was performed using 920 nm illumination. Images were acquired with a 2.3 × 2.3 μm pixel size, and 5 μm plane spacing. 8-10 optical planes were acquired over a depth of 50 μm in total. To image the entire brain, images were acquired as tilesand stitched using StitchIt (https://doi.org/10.5281/zenodo.3941901). After each mosaic tile was imaged at all ten optical planes, the microtome automatically cut a 50 μm slice, enabling imaging of the subsequent portions of the sample and resulted in full 3D imaging of entire brains. All images were saved as a series of 2D TIFF files.

Images were registered to the Allen Mouse Brain Common Coordinate Framework (Wang et al., 2020) using the software brainreg (Tyson et al., 2022) based on the aMAP algorithm (Niedworok et al., 2016). All atlas data were provided by the BrainGlobe Atlas API (Claudi et al., 2021). For registration, the sample image data was initially down sampled to the voxel spacing of the atlas used and reoriented to align with the atlas orientation using bg-space (https://doi.org/10.5281/zenodo.4552537). The 10 μm atlas was used for cell detection and mapping. To manually segment structures within the brain (i.e. barrels in SSp-bfd) as well as analyse and summarize specific tissue volumes and viral injection sites, the software brainreg-segment (Tyson et al., 2022) was used. Automated cell detection and deep learning based cell classification was performed using the cellfinder software (Tyson et al., 2021) and cross-validated with manual annotation. All analysis in this manuscript was performed in atlas space (Wang et al., 2020).

Figures showing detected cells in 3D atlas space were generated using the brainrender software (Claudi et al., 2021) and custom scripts written in Python 3.9. The total number of detected brain cells varied from animal to animal. Therefore, cell numbers in different brain areas are reported as a fraction of the total number of cells per animal detected within the brain areas given in the specific diagrams. For comparison of the laminar distribution of cells within different brain areas, values were normalized to the total number of cells detected in each area. If not stated otherwise, diagrams always present the fraction of detected neurons in the hemisphere ipsilateral to the injection site. Dorsal views of cortical areas are maximum projections along the dorso-ventral axis.

To investigate the spatial distribution of cells in areas of interest (VISp, and L6 and L2/3 of SSp-bfd) we used principal component analysis (PCA). This aid us in determining the main directions over which the data were dispersed and enabling us to section each area spatially in a non-arbitrary manner. For analysis, we used the first two principal components (2D vectors). Mathematically, the first principal component is the direction in space along which projections have the largest variance. The second principal component is the direction which maximizes variance among all directions orthogonal to the first. To compute these components for each area, we first projected the cell position coordinates (3D) to the 2D spatial space of interest, pooled them from all corresponding mice, and then applied PCA. The pre-PCA projection to 2D space enabled direct mapping of the principal components to the spatial axes of our data (i.e. anatomical axes) thereby rendering them more intuitive. Of note, in our preliminary analysis, we obtained very similar results when computing PCA per mouse and then averaging over mice for either of principal components. The variances explained by the first and second principal components for our data were: [80%, 20%] for L2/3 of SSp-bfd, [65%, 35%] for L6 of SSp-bfd, and [61%, 39%] for VISp. The normalized amplitude of each plotted principle component shows its explained variance relative to the other one. For this analysis, we used the *PCA* module of *Scikit-learn* library in Python 3.9.

For sectioning based on 1^st^ principal component, using standard linear algebra techniques and the fact that the 2^nd^ principal component is orthogonal to the 1^st^ one, we computed a set of lines (i.e. sections borders) having a slope equal to that of 2^nd^ principal component, and perpendicular to the virtual expansion of 1^st^ principal component (in 2D space, with same slope). Thus, sectioning lines are parallel to each other and parallel to the 2^nd^ principal component. We set the distance between each two lines equal to 200 pixels, which was implemented by adjusting their y-intercepts. The sectioning based on 2^nd^ principal component was performed similarly and only used in combination with that of 1^st^ principal component for parcellation of horizontal projections of VISp neurons.

To compute the smoothed cell density map of each area we evaluated a Gaussian kernel density on a 2D regular grid with uniformly spaced x-coordinates and y-coordinates (the anatomical coordinates) in the intervals limited to the maximum and minimum values of their coordinates. By sufficiently padding these limits (or borders) we also relaxed the potential edge effects, and cut them off after applying the Gaussian kernel. We applied these steps to each mouse separately and also determined the position of its maximum density (2D). To compute an overall single map for each area of interest, we averaged over corresponding mice. To better visualize the variation and the relative gradient of cell densities in 2D space we also computed the contour lines of this map. The points on each contour line have the same cell density, and the gradient of the cell density is always perpendicular to the contour lines where the closer distance between the lines reflects a larger gradient (i.e. steeper variation in cell density). All these steps were performed in Python 3.9. To create the spatial grid we used *mgrid* module of *numpy* library with a step length of 58 (i.e. 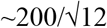) pixels. To apply the Gaussian kernel density we used *kde* function in *stats* module of *scipy* library and enabled its automatic bandwidth determination method called *scott*. To plot the density maps and contour lines we used *pcolormesh* and *contour* functions, respectively, in *pyplot* module of *matplotlib* library. When using *contour* function, we enabled the option for automatic selection of the number and position of the lines.

To compute the fraction of cells in each barrel column, we first selected the barrels as region of interests in Fiji (https://fiji.sc/) based on the reconstructed entire barrel field in layer 4, and then imported them to Matlab 2020a (MathWorks) and created a mask for each barrel using *ReadImageJROI* and *ROIs2Regions* functions (https://github.com/DylanMuir/ReadImageJROI), respectively. Next, for each mouse we counted the number of cells located in each barrel, separately for layer 6a and layer 2/3. In order to have the same scale over these layers, we computed the fraction of cells per barrel by dividing its cell count by the total number of cells in both layers. To investigate which barrel has a cell count beyond the chance we performed a randomization test. To do this for each layer, we computed *fr_b_* = (#Cells_*b*_ / #Cells_tot_)×(#Area_tot_ / #Area*_b_*) as the relative fraction index of each barrel while accounting for different barrel sizes; #Cells_*b*_ and #Area_*b*_, (resp. #Area_*b*_ and #Area_tot_) are the cell count and number of pixels of barrel *b* (resp. of whole depicted barrel field). We then uniformly shuffled the position of all cells in each layer over the whole barrel filed and re-computed the relative fraction index using the same formula and called it 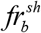. We repeated this step for 2500 times, yielding a distribution of 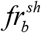. Finally, to assess how likely is to observe *fr_b_* of each barrel in the randomized data, we adapted one-tailed permutation test of Cohen thereby accounting for multiple comparison problem; the significance level was set to 0.001.

To assign visual space coordinates to postsynaptic neurons in VISp (horizontal projection), we first parceled VISp into 31 subareas, based on sectioning along the 1^st^ and 2^nd^ PCs of the postsynaptic neuron distribution (see above). After determining the average fraction of neurons within each parcel we generated a color coded input map of VISp within the areal border of VISp from the CCFv3. This border was then aligned with the mean field sign borders of VISp containing elevation and azimuth contour plots (Zhuang et al., 2017). Subsequently, elevation and azimuth coordinates were assigned for each parcel of VISp to estimate the extent of the modulated visual field based on the fraction of postsynaptic neurons.

### Optophysiology

The cutting solution for in vitro experiments contained 85 mM NaCl, 75 mM sucrose, 2.5 KCl, 24 mM glucose, 1.25 mM NaH2PO4, 4 mM MgCl2, 0.5 mM CaCl2 and 24 mM NaHCO3 (310-325 mOsm, bubbled with 95% (vol/vol) O2, 5% (vol/vol) CO2). Artificial cerebrospinal fluid (ACSF) contained 127 mM NaCl, 2.5 mM KCl, 26 mM NaHCO3, 2 mM CaCl2, 2 mM MgCl2, 1.25 mM NaH2PO4 and 10 mM glucose (305-315 mOsm, bubbled with 95% (vol/vol) O2, 5% (vol/vol) CO2). Caesium-based internal solution contained 122 mM CsMeSO4, 4 mM MgCl2, 10 mM HEPES, 4 mM Na-ATP, 0.4 mM Na-GTP, 3 mM Na-L-ascorbate, 10 mM Na-phosphocreatine, 0.2 mM EGTA, 5 mM QX 314, and 0.03 mM Alexa 594 (pH 7.25, 295-300 mOsm). K-based internal solution contained 126 mM K-gluconate, 4 mM KCl, 10 mM HEPES, 4 mM Mg-ATP, 0.3 mM Na-GTP, 10 mM Na-phosphocreatine, 0.3% (wt/vol) Neurobiotin tracer (pH 7.25, 295-300 mOsm).

Acute brain slices were obtained on the day of in vitro experiments, as previously described (Weiler et al., 2018). In brief, mice were deeply anesthetized with isoflurane (~2%) in a sealed container and rapidly decapitated. Coronal sections of VISp (320 μm) were cut in ice cold carbogenated cutting solution using a vibratome (VT1200S, Leica). Slices were incubated in cutting solution in a submerged chamber at 34°C for at least 45 min and then transferred to ACSF in a light-shielded submerged chamber at room temperature (21-25°C) until used for recordings. The expression patterns of ChR2-EGFP as well as td-tomato within VISp and SSp-bfd were screened using fluorescence detection googles (Dual Flourescent Protein Flaslight, Nightsea) with different excitation light (cyan and green) and filters during the slice preparation. Only slices with visibly sufficient transduction of ChR2-EGFP were considered for experiments. Brain slices were used for up to 6 h. A single brain slice was mounted on a poly-D-lysine coated coverslip and then transferred to the recording chamber of the microscope while keeping track of the rostro-caudal orientation of the slice. All recordings were performed at room temperature (20-25 °C).

Brain slice visualization and recordings were performed on an upright microscope (Scientifica Slice Scope Pro 600) using infrared Differential Interference Contrast (DIC) with a low magnification objective (4x objective lens) and images were acquired by a high-resolution digital CCD camera.

VISp was identified using morphological landmarks and the presence of fluorescent axons. Whole cell recordings were performed at high magnification using a 40× water-immersion objective. Targeted cell bodies were at least 50 μm below the slice surface. Borosilicate glass patch pipettes (resistance of 4-5 MΩ) were filled with a Cs-based internal solution for measuring excitatory and inhibitory postsynaptic currents in the same cell (EPSC: voltage clamp at −70 mV, IPSC: voltage clamp at 0 mV). K-based internal solution was used when recording EPSC, postsynaptic potentials (EPSs) and sub- and suprathreshold electrophysiological properties. Basic electrophysiological properties were examined in current-clamp mode with 1 s long hyper- and depolarizing current injections. Once stable recordings with good access resistance were obtained (< 30 MΩ), recordings were started.

Data were acquired with Multiclamp 700 B amplifiers (Axon Instruments). Voltage clamp recordings were filtered at 10 kHz and digitized at 20 kHz. The software program wavesurfer (https://wavesurfer.janelia.org/index.html) in MATLAB 2019b (Mathworks) was used for hardware control and data acquisition.

For ChR2 photostimulation, LED light was generated using a light emitting diode (LED) (470 nm) and controlled by a CoolLED pE-300ultra system (CoolLED). Collimated light was delivered into the brain tissue through the 40x objective. For the input mapping experiments, different trains of photostimuli were delivered: (1) 5 pulses with 10 Hz and 100 ms duration of each pulse. (2) 25 pulses with 5 Hz and 10 ms duration. (3) 50 pulses with 10 Hz and 10 ms duration. The laser intensity for each pulse was set to ~ 5mW/mm2. In a subset of recordings, tetrodotoxin (TTX, 1 μM, Merck) and 4-aminopyridine (4-AP) (100 μM, Merck) was bath perfused to isolate direct monosynaptic inputs during photostimulation.

Intrinsic electrophysiological parameters were extracted using the PANDORA Toolbox (Gunay et al., 2009) and custom-written software in MATLAB 2019-2022. The suprathreshold single spike parameters were measured using the first spike evoked by current injection (at Rheobase). For photostimulation experiments, light evoked PSCs as well as PSPs were considered non-zero if their amplitudes were larger than 7 times the standard deviation of a 100 ms baseline directly before stimulus onset. Additionally, suprathreshold responses were only included if they occurred at least twice in photostimulation trains (2) and (3) (see above). The inflection points of the EPSCs and IPSCs were defined as the onsets and used to calculate the onset latencies with custom-written software in MATLAB 2019-2022.

### Neuronal network modeling

To gain insights into the underlying network mechanism of active VISp suppression via SSp-bfd activation, we used computational modeling and mathematical stability analysis. To this end, we formulated neural dynamics of each cortical area using a well-established, canonical cortical recurrent neural network (RNN) model of mean firing activity rates of two spatially localized, homogeneous glutamatergic and GABAergic cells (here, pyramidal (PN) and fast-spiking (FS) populations; according) (Murphy and Miller, 2009; Tsodyks et al., 1998; Wilson and Cowan, 1972). This Wilson-Cowan-type model and its extensions have the advantage of being biophysically interpretable, mathematically accessible and amenable to replicate the behavior of spiking models, and has been used extensively in previous studies to explain the behavior and underlying mechanisms of cortical networks (Murphy and Miller, 2009; Ozeki et al., 2009; Rahmati et al., 2017; Tsodyks et al., 1998; Tsodyks et al., 1997). We connected, extended and parameterized the two RNNs (i.e. two pairs of PN-FS populations) based on our present data and previous studies (see below). In the following, we first formulate the model dynamics mathematically and extend it to account for our observed cross-modal input, outline the rationale of model parameterization in the present work, describe the applied stability analysis including the operating regimes, and finally define the quantities which we extracted from our modeling results. For brevity, hereafter, in the mathematical expressions we refer to VISp and SSp-bfd regions as V and S (mainly as superscripts like RNN^V^ and RNN^S^), and to PN and FS as P and F.

#### Model description and extension

The equations governing the mean-field dynamics of the RNN^S^ over time are (dots denote the time derivatives) (Tsodyks et al., 1997; Wilson and Cowan, 1972):

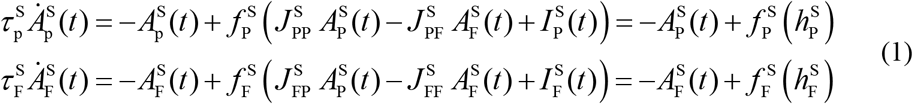

where 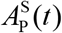 and 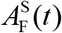 are the average activity rates (in Hz) of PN and FS populations which can be properly scaled to represent locally the average recorded activities in these populations, 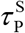 and 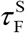 are the time-constants of PN and FS populations’ firing activities to approach their steady states, 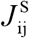 (i and j ∈{P,F}) are the average synaptic weights of recurrent (i=j) or feedback (i ≠ j; j→i) connections within SSp-bfd, and 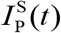 and 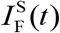 are the external inputs received by SSp-bfd’s PC and FS populations from other brain regions or via stimulation. All synaptic weights have positive values.

The RNN^V^ population dynamics can be described similarly to Eq. 1. However, here, we extend this regional model in order to account for the observed cross-modal projection and its different postsynaptic effects (depolarizing vs. excitatory), as observed in our data:

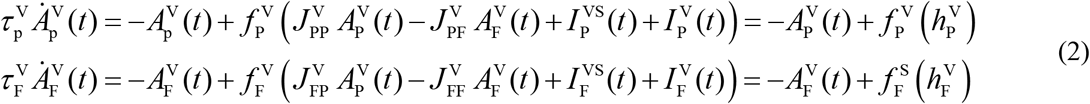

where 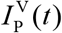 and 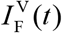 are the external inputs received by VISp’s populations from other brain regions than SSp-bfd or via stimulation, and 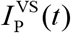 and 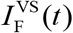 are the glutamatergic synaptic-inputs from SSp-bfd’s PN population onto VISp’s PN and FS populations with corresponding average synaptic weights 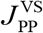 and 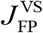, respectively, which we modeled as:

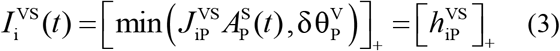

where []_+_ is a threshold-linear function formulated as 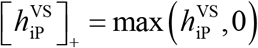, and we defined δ as a positive constant controlling the depolarizing or excitatory action of the cross-modal input (see next section for its parameterization). In both Eqs. 1 and 2, the transformation from the summed input to each population 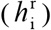 to an activity output (in Hz) is governed by the response function 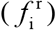 defined as(Rahmati et al., 2017; Tsodyks et al., 1998)

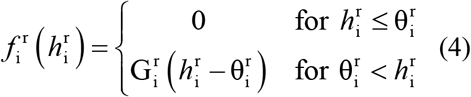

where r ∈{V,S}, 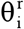 is the population activity-threshold, and 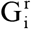 is the linear input-output gain. The effective weights of the synaptic projection in the active mode 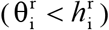 can be defined as 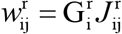 and 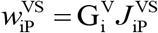, where again i and j refer to post- and presynaptic populations, respectively. Similarly, we define the effective values of external inputs (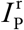 and 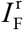) as 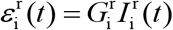.

#### Parameterization

To set the parameter values of each RNN we mainly followed previous studies (Murphy and Miller, 2009; Ozeki et al., 2009; Rahmati et al., 2017; Tsodyks et al., 1997), and used our experimental data to qualitatively constrain the parameterization of our cross-modal network model (Eqs. 1–4), as follows. (**i**) Since EPSCs mediated by SSp-bfd onto VISp’s PN and FS populations exhibited similar amplitudes, we considered 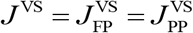, thus 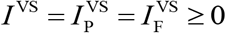. (**ii**) Since these EPSC amplitudes onto VISp’s PN did not change in presence of TTX+4-AP, we considered 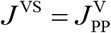. (**iii**) Since the evoked IPSCs (mediated locally by FS INs within VISp, following the activation of axonal fibers axonal fibers from SSp-bfd) onto VISp’s PNs were around two times bigger than the evoked EPSCs, we considered 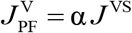 (similarly, 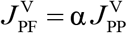) with α = 2; where, α is analogous to 1 over I/E ratio. (**iv**) Since the VISp’s FS INs, as compared to PNs, exhibited higher intrinsic excitability (smaller difference between RMP and AP-threshold and less negative RMP) we considered 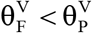, where in accordance to our data we set 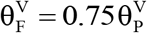 with 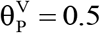. (**v**) Since FS INs exhibited a much higher gain than PNs, we considered 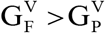 and set their default values proportionally, in accordance to our data, as 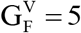 and 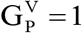. (**vi**) Since the glutamatergic projection from SSp-bfd onto VISp’ PNs was depolarizing rather than excitatory, we set an upper-limit on *I*^VS^ using Eq. 3, defined as 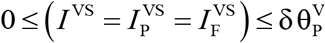. Here, we set δ = 1 so that the isolated effect of *I*^VS^ on PNs will be depolarizing only; i.e. to be smaller than the population activity-threshold of PN population, 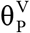 (see Eq. 4). This modeling enables *I*^VS^ to act as both excitatory/depolarizing on VISp’ FS INs since 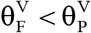 (see iv). Intuitively, whereas the strongest action of *I*^VS^ alone on the PN population is just to bring it closer to the active mode, the same *I*^VS^ can even enable firing of FS population (if 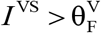). Of note, in our preliminary analysis we obtained very similar results by setting e.g. δ = 1.1 (dominantly depolarizing and moderately excitatory). (**vii**) Finally, to provide a clearer representation of our main modeling results about the suppression phenomenon as well as to avoid fine-tuning of parameter values, we made, without losing the generality, the following two assumptions. First, we forwent the potential regional differences in the parameter values as we built our cross-modal network model based on the canonical cortical RNN model used to model either of regions (see above). Nonetheless, our preliminary analysis showed that this assumption could be also neglected since the main role of RNN^s^ in this context is to drive the cross-modal input and thus, could be modeled by considering only its cross-modal inputs to RNN^v^ (e.g. like a stimulus). Second, we assumed that the intra-areal glutamatergic projections (PN→PN and PN→FS) and GABAergic projections (FS→FS and FS→PN) have the same effective weights, separately (see also e.g. (Murphy and Miller, 2009; Rahmati et al., 2017)): 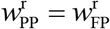 and 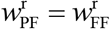; although, note that the (non-effective) synaptic weight 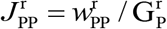 (resp. 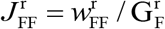) is not necessarily equal to 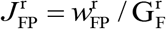 (resp. 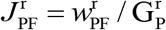) since 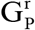 and 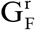 can have different values (as in the present work). Of note, in our preliminary results, we observed that our main findings remained largely intact even without these assumptions. Despite, considering these assumptions enabled us to specifically focus on and encapsulate the corresponding effect of gains on processing of cross-modal input in VISp. Accordingly, we considered 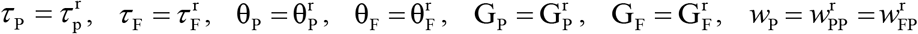, and 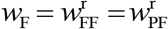, where r ∈ {V,S}. Consistent to previous studies (Murphy and Miller, 2009; Ozeki et al., 2009; Rahmati et al., 2017), we set *τ*_p_ = 60ms, *τ*_F_ = 12ms, and *w*_P_ = 2. Then, according to our aforementioned constraints we obtain: 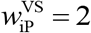, and *w*_F_ = 4. The rest of parameter values can be set according to the aforementioned description. In addition, note that when investigating the effect of VISp’s population-gains on the suppression we varied 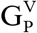 and 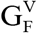 values independently from SSp-bfd’s firing gains (i.e. 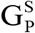 and 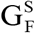 were kept at their default values 1 and 5, respectively; see point v). This allowed us to specifically assess the significance of firing gains of VISp’s populations on the suppression phenomenon independently from the effect of SSp-bfd’s gains, as this strategy keeps the amount of cross-modal input the same. In our simulations, we set the effective values of external inputs to 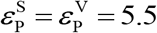 and 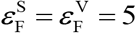 with the onset and offset times illustrated in **Figure 7B**. For each region, the inputs to PN and FS populations were applied (and ceased) simultaneously.

#### Phase plane and fixed point

To visualize the stability behavior of RNN^V^ model, we used the phase plane analysis based on the activity rates: A_FS_-A_PN_-plane. The A_FS_-A_PN_-plane sketch includes the curves of the A_PN_ and A_FS_-nullclines representing sets of points for which 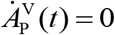 and 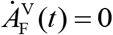. Any intersection of these nullclines is called a fixed point (FP or steady state), with the stability needed to be determined (see the following text). The FP represents the steady state levels of RNN^V^ population activities (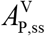 and 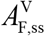; constant values) in presence of a sufficiently long-lasting external (visual stimulation) and/or cross-modal inputs. The two FPs in our results are approached during visual stimulation (v; FP_v_) or visual+tactile stimulations (v + w; FP_v+w_) at the corresponding steady state of the network. Note that at the FP(s) the cross-modal input can be considered to have approached its steady state value: 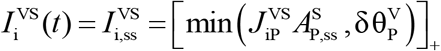. Accordingly, to plot the A_FS_-A_PN_-plane we can substitute the 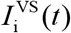 with its steady state value 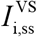, thus only solve the 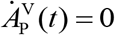 and 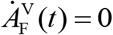 equations; i.e. decouple Eqs. 1 and 2. Different levels of 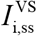 can mainly push the nullclines up or down in the A_FS_-A_PN_-plane, without changing their slopes. To determine the stability of each FP we applied the linear stability analysis to RNN^V^ system of equations in Eq. 2 while considering 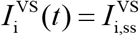: We investigated whether all eigenvalues of the corresponding Jacobian matrix have strictly negative real parts (if so, the FP is stable), or whether at least one eigenvalue with a positive real part exists (if so, the FP is unstable). The details about plotting the phase planes and stability analysis have been described in previous studies (Ozeki et al., 2009; Rahmati et al., 2017; Tsodyks et al., 1997).

#### Operating regimes

The stable operating regimes of a RNN at a FP can be classified as an inhibition-stabilized network (ISN) vs. a Non-ISN (Ozeki et al., 2009; Rahmati et al., 2017; Tsodyks et al., 1997). To apply this theoretical classification to the RNN^V^ model we applied the previously described analytical and numerical techniques (for details see (Ozeki et al., 2009; Rahmati et al., 2017)) to Eq. 1 while considering 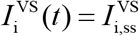 (see previous paragraph). In brief, to discriminate between these two regimes three criteria were defined: (A) Excitatory instability: For the inhibitory activity (here of FS-population) rate fixed at the FP, the recurrent excitation is strong enough to render the PN-population intrinsically unstable. (B) Excitatory stability: In contrast to (A), the PN-population is stable per se, i.e. even with a feedback inhibition fixed at its level at the FP. (C) Overall stability: The dynamic feedback inhibition to the PN-population is strong enough to stabilize the whole network activity. At a FP, a network operating under the (A) and (C) criteria is an ISN, while a network operating under the (B) and (C) criteria is a Non-ISN. A network, which is neither ISN nor Non-ISN at the FP, operates under an unstable regime. In the A_FS_-A_PN_-plane, (A) and (B) render the slope of A_PN_-nullcline positive and negative, respectively. Under (C), the slope of A_FS_-nullcline (positive) is bigger than the slope of A_PN_-nullcline at the FP. RNN^V^ operates as an ISN at the FPs shown in our results, since both A_PN_- and A_FS_- nullclines have positive slopes and A_FS_-nullcline’s slope is steeper (Ozeki et al., 2009; Rahmati et al., 2017).

#### Simulations

All modeling simulations in this paper have been implemented as Wolfram Mathematica 13 and Matlab 2020a (MathWorks) code. For network simulations, we set the integration time-step size to 0.0001 s. The initial values of cross-model network’s variables were set to 0, and it was simulated for 15 seconds; i.e. 5 seconds under each external input configuration, thereby assuring the convergence of firing activities to their steady states under each configuration. For RNN^V^, the input order was: initialization (no input), v only, and v + w. Accordingly, the RNN^S^ was simulated 10 seconds without input, and then the tactile-resembling input (w) was added.

#### Additional quantities

We quantified the change in RNN^V^ activity under the presence of cross-modal input as the difference between the steady-state value of its sum activity during v + w stimulation 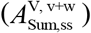 and that during v-only stimulation 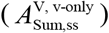, i.e. 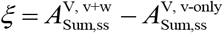; *ξ* < 0: suppression, *ξ* > 0: no suppression (e.g. facilitation), *ξ* = 0: unchanged. Both absolute and relative amplification parameters quantify the strength of the transient increase in RNN^V^’s PN activity (before its overall suppression) upon the onset of tactile stimulation. We formulated these parameters as 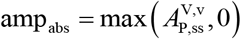 and 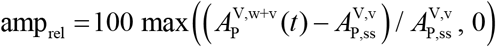, respectively, where 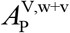 and 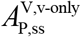 are the running value and the steady state value of PN activity during the corresponding stimulations (see their superscripts). We showed these parameters in dB, while considering the values of 0 (i.e. no amplification) as undefined. Finally, to quantify the effective time of RNN^V^’s FS-mediated inhibition to gain up, we subtracted the peak-time of 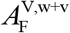 from that of the transient amplification in RNN^V^’s PN activity 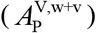 under v + w stimulation.

### Statistical analysis

Details of all *n* numbers and statistical analysis are provided in all figure captions. The required sample sizes were estimated based on literature and our past experience performing similar experiments. Significance level was typically set as p<0.05 if not stated otherwise. Statistical analyses were performed using GraphPad Prism v.9, MATLAB 2019-2022 and Python 3.9.

## 6 Acknowledgements

We thank Michael Richter and Elisabeth Meier for excellent for technical support and technical assistance. The authors are further grateful to the support staff of the Neurobiological Research Facility at Sainsbury Wellcome Centre.

This project is supported by the Interdisciplinary Centre for Clinical Research (IZKF; Advance medical scientist - Program 11) and by funding from the Foundation “Else Kröner-Fresenius-Stiftung” within the Else Kröner Graduate School for Medical Students “Jena School for Aging Medicine” (JSAM). T.W.M. and S.W. are funded by The Wellcome Trust (214333/Z/18/Z; 090843/F/09/Z) and Humboldt Foundation (S.W.). K.H. is funded by the German Research Foundation (HO 2156/5-1, HO 2156/6-1). A.S. and C.F. are funded by the German Federal Ministry for Economic Affairs and Climate Action (BMWK) within the Promotion of Joint Industrial Research Programme (IGF) due to a decision of the German Bundestag as part of the research project (IGF 22462 BR) by the Association for Research in Precision Mechanics, Optics and Medical Technology (F.O.M.) under the auspices of the German Federation of Industrial Research Associations (AiF). C.G. is funded by the German Research Foundation (FOR3004; GE 2519/8-1; GE 2519/9-1).

## 8 Author contributions

S.W. conceived the project, performed tracing and electrophysiological experiments, analyzed electrophysiological experiments, interpreted all data and wrote the manuscript. V.R. contributed to the analysis of tracing data, performed network modeling and wrote the manuscript. J.W. generated, animated and analyzed the model of the mouse whisker array. M.I. performed imaging experiments and immunohistological stainings. A.W.S. contributed to the generation of the model of the mouse whisker array. C.F., C.G. and O.W.W. provided resources and interpreted data. M.H. interpreted data and wrote the manuscript. J.B. supervised imaging experiments and imaging data analysis, interpreted data and wrote the manuscript. T.M. and K.H. supervised the project and wrote the manuscript. M.T. designed the experiments, analyzed and interpreted the majority of data, supervised the project and wrote the first draft of the manuscript.

