## Supplemental information for "A primary sensory cortical interareal feedforward inhibitory circuit for tacto-visual integration"

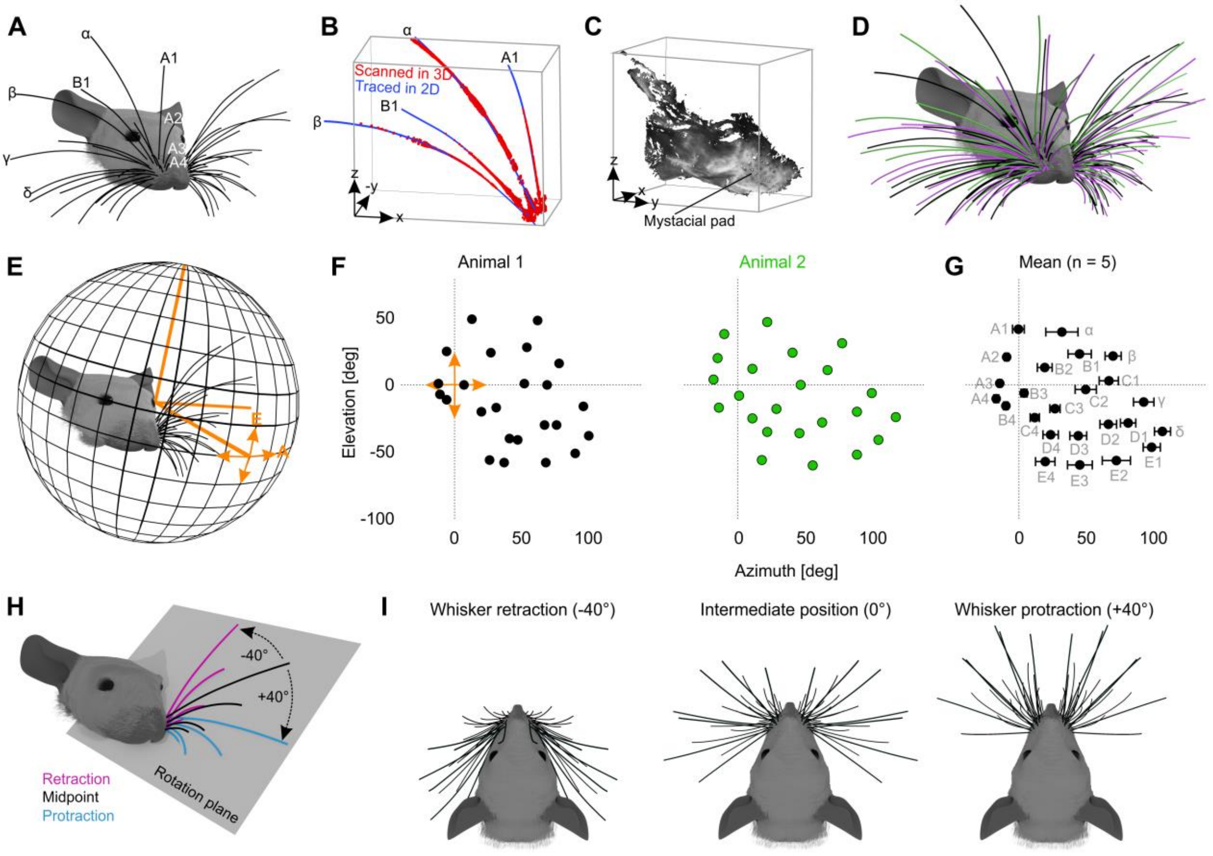


**Supplemental Figure 1: Reconstruction of the mouse whisker array.** (A) Representative 3D model of the mouse whisker array aligned with a realistic 3D model of the mouse head (Bolanos et al., 2021). Letters indicate the standard nomenclature for the labeled whiskers. (B) After their 3D-reconstruction all whiskers were plucked, traced in 2D and then aligned to their corresponding 3D point clouds. (C) 3D point cloud of the mouse head including the mystacial whisker pad (obtained after whisker plucking). (D) Morphologically accurate model of the mouse head together with 3D-reconstructed superimposed whiskers from three different exemplary mice. (E) An eye-centered spherical coordinate system was used to determine the position of each whisker tip in elevation (E) and azimuth (A) with respect to the eye. (F) Representative elevation and azimuth coordinates of all 24 large whisker tips with respect to the left eye (from two exemplary mice). (G) Average elevation and azimuth coordinates of all whisker tips. Centroids represent their average position ± s.e.m. (n=5). Letters indicate the standard nomenclature of the 24 large whiskers. (H) For simulation of whisker retraction and protraction, whiskers of each row were rotated around their base point to -40° or +40° along a plane fitted through their base points and tips. (I) Representative examples of whisker retraction (-40°), intermediate position (0°) and protraction (+40°).

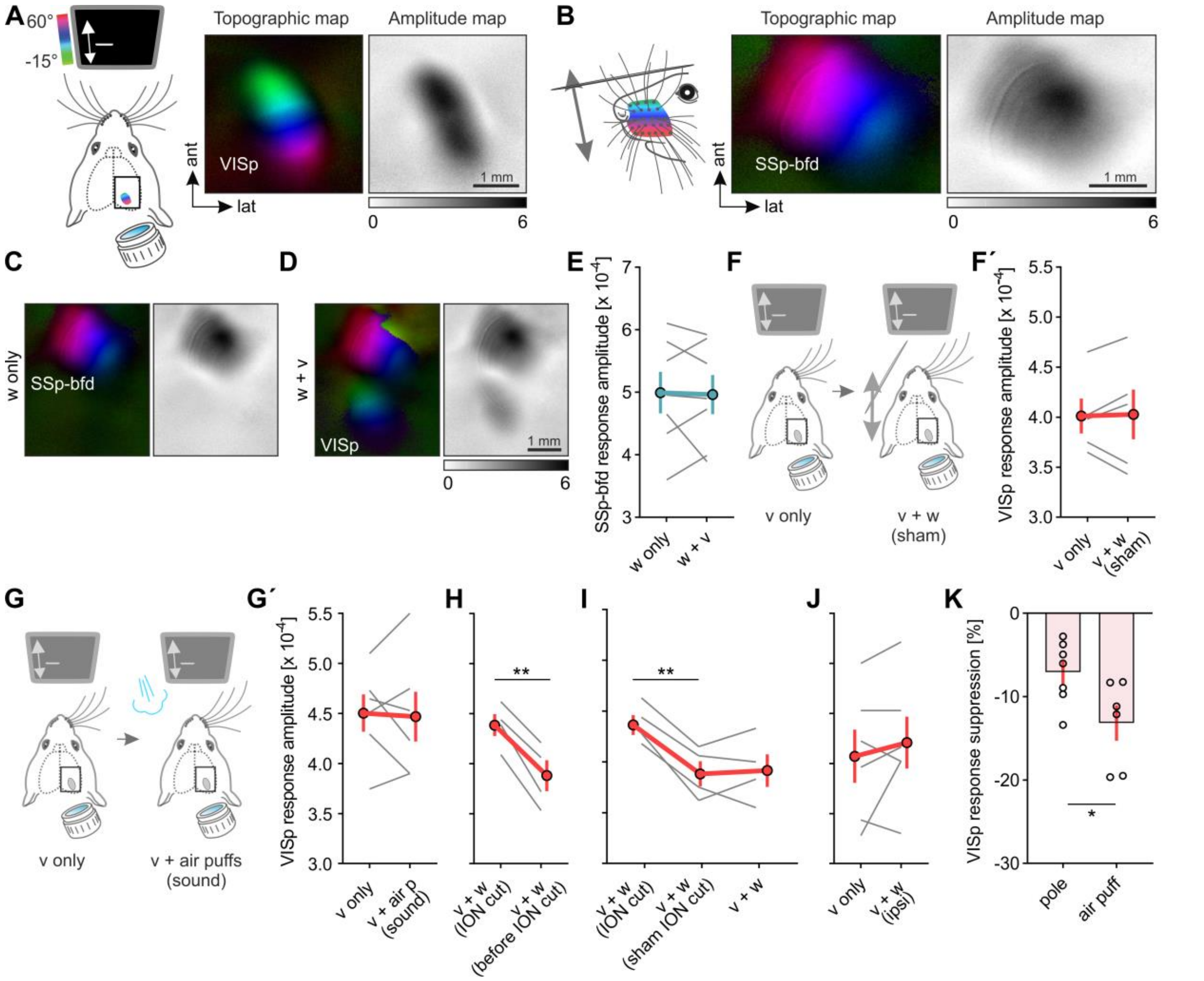


**Supplemental Figure 2: Whisker stimulation suppresses visually driven activity in VISp in vivo. Related to Fig. 2.** (A, B) Schematics of unimodal visual or whisker stimulation procedures together with the obtained topographic and amplitude maps of VISp and SSp-bfd. (C, D) Cortical topographic and amplitude maps evoked after whisker stimulation alone (w only) and bimodal whisker and visual stimulation (w+v). (E) Quantification of SSp-bfd response amplitudes under w only and w+v conditions (n=7). (F) Schematic of unimodal visual and sham whisker stimulation after whisker removal. (F´) Quantification of VISp response amplitudes (n=5, p=0.8744; paired t-test). (G) Schematic of stimulation procedures to test for the influence of the sounds created by the air puffs on visually driven VISp activity. In the bimodal stimulation condition, air puffs were directed to the same space as for whisker stimulation, however, after the whiskers were removed. (G´) Quantification of VISp response amplitudes evoked under the stimulus conditions depicted in g (n=6, p=0.8173; paired t-test). (H) Quantification of VISp response amplitudes evoked under bimodal stimulation (v+w; w: air puffs) before and after ION cut (n= 4, p=0.0049; paired t-test). (I) Quantification of VISp responses evoked under bimodal stimulation conditions (v+w; w: air puffs) before and after a sham ION cut and after finally cutting the ION (n= 4, p=0.0368, p=1; paired t-test followed by Bonferroni correction). Sham ION cut was performed by opening the skin above the ION and touching the nerve gently with fine forceps. (J) Quantification of VISp responses evoked under v only and v+w stimulation conditions (n=6, p=0.3742; paired t-test). Here, whiskers ipsilateral to the recorded hemisphere were stimulated using air puffs. (K) Percentage suppression of visually evoked VISp activity by simultaneous whisker stimulation with either the pole or air puffs (n=7, n=6, p=0.03; unpaired t-test). In E-J: Solid grey lines represent paired measurements of individual mice and red lines represent means ± s.e.m. In K: Circles indicate measurements of individual mice and bars represents means ± s.e.m. *p<0.05, **p<0.01, ***p<0.01

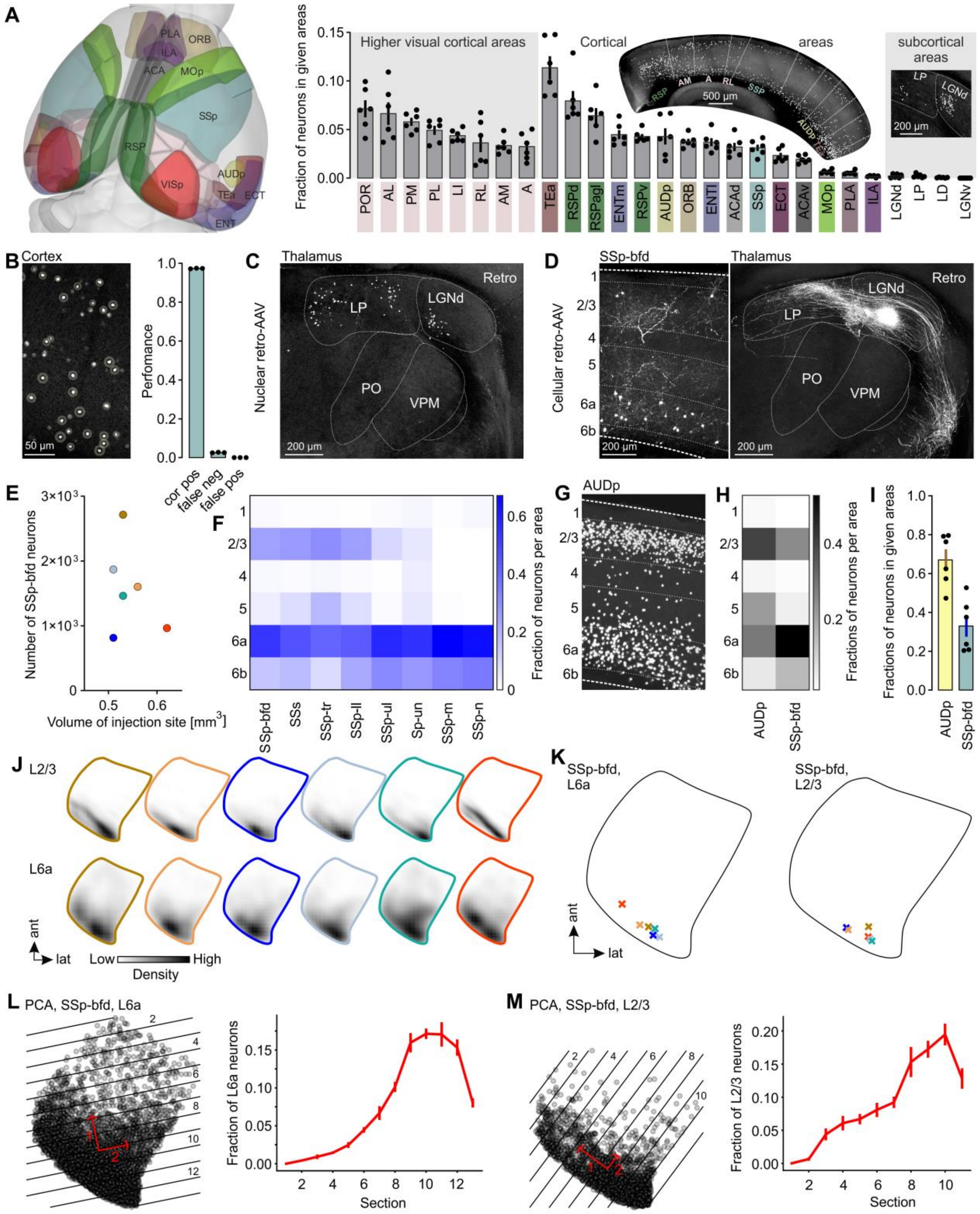


**Supplemental Figure 3: Analysis of the distribution of neurons projecting to VISp.** (A) Left, 3D rendered mouse brain with colored cortical brain areas containing VISp projecting neurons. Right, Fraction of projection neurons in different ipsilateral brain areas. Black circles indicate fractional cell counts of individual mice (n=6). Bars represent means ± s.e.m. (B) Left, retrogradely labeled neurons in the mouse cortex. Neurons correctly detected by cellfinder (Tyson et al., 2021) are marked by a ring. Right, performance of cellfinder in detecting labeled neurons correctly. Black circles indicate fractional cell counts of individual mice (n=3). Bars represent means. (C, D) Example images showing neurons in thalamic areas and SSp-bfd retrogradely labeled by either the nuclear or the cellular retro-AAV (AAV-EF1a-H2B-EGFP, rAAV2-retro.CAG.GFP) after their injection into VISp. Numbers in (D) indicate cortical layers. (E) The absolute number of detected projection neurons in SSp-bfd from 6 different mice was plotted against the volume of the injection site in VISp (r=-0.3115, p=0.5478). (F) Color-coded average fraction of projection neurons in cortical layers across somatosensory cortical areas (n=6). Numbers on the left indicate cortical layers. Values are normalized to the maximum within each area. (G) Representative coronal section showing retrogradely labeled neurons in AUDp. Numbers indicate cortical layers. (H) Grey-scale-coded average fraction of projection neurons in cortical layers of AUDp and SSp-bfd in the same mice (n=6). Numbers indicate cortical layers. Values are normalized to the max within each area. (I) Fraction of projection neurons in AUDp and SSp-bfd. Black circles indicate fractional cell counts of individual mice (n=6). Bars represent means ± s.e.m. (J) Density maps of projection neurons in L6 and L2/3 of SSp-bfd (horizontal projection) of 6 individual mice. Border colors indicate the corresponding injection sites in VISp (see Figure 3I). (K) Horizontal projections of L6 and L2/3 of SSp-bfd. Crosses indicate the location of the maximum of density of the 6 different projections neuron populations. Colors relate to the different injection sites in VISp (see Figure 3I). (L, M) Left, horizontal projections of L6 and L2/3 of SSp-bfd. Grey circles indicate projections neurons labeled by all 6 injection sites in VISp. Red arrows indicate the first and second principal components determining the direction of the spatial dispersion of projection neurons. Black lines, section borders orientated vertically to the first principle component. Right, the average fraction of projection neurons in all sections of L6 and L2/3 (n=6). PCA: Principal component analysis.

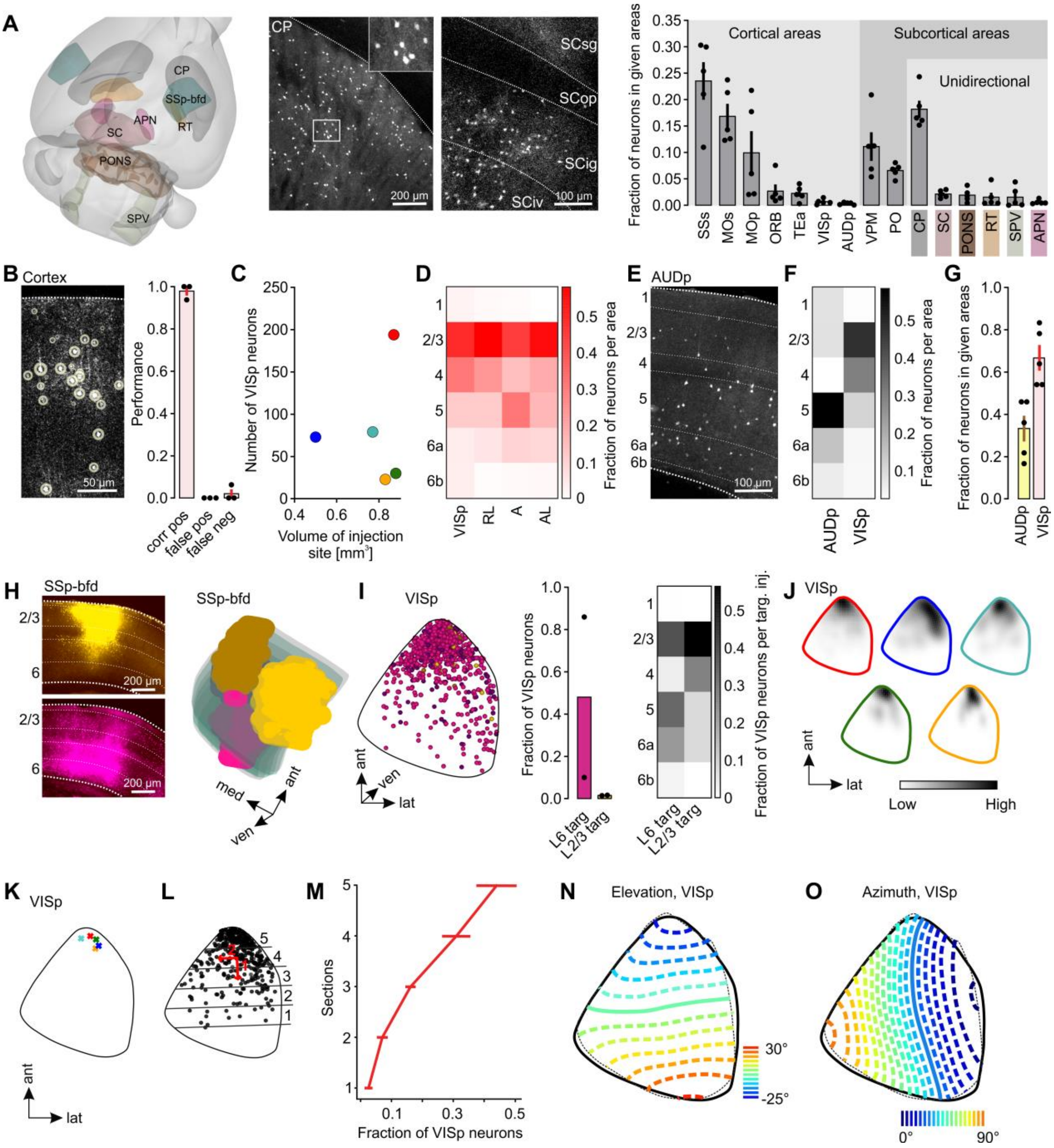


**Supplemental Figure 4: Related to Fig. 5.** (A) Left, 3D-rendered mouse brain with colored subcortical brain areas uni-directionally targeted by SSp-bfd. Middle: Representative coronal images of postsynaptic neurons in the caudoputamen (CP) and superior colliculus (SC). Right, fraction of postsynaptic neurons in different cortical and subcortical brain areas. Black dots indicate fractional cell counts of individual mice (n=5). Bars indicate means ± s.e.m. (B) Left, anterogradely labeled neurons in the mouse cortex. Neurons correctly detected by the cellfinder (Tyson et al., 2021) software are marked by a ring. Right, performance of cellfinder in detecting labeled neurons correctly (n=3 mice). (C) The absolute number of detected postsynaptic neurons in VISp from 5 different mice was plotted against the volume of the corresponding injection site in SSp-bfd (r=0.1028, p=0.8694). (D) Color-coded average fraction of postsynaptic neurons across cortical layers in visual cortex areas (n=5). Numbers on the left indicate cortical layers. Values are normalized to the max within each area. (E) Representative coronal image showing postsynaptic neurons in AUDp (white). (F) Grey-scale-coded average fraction of postsynaptic neurons across cortical layers of AUDp in comparison to VISp (n=5). Numbers indicate cortical layers. Values are normalized to the maximum within each area. (G) Fraction of postsynaptic neurons in AUDp and VISp. Black dots indicate fractional cell counts of individual mice (n=5). Bars indicate means ± s.e.m. (H) Left, coronal images showing representative injection sites targeted to L2/3 and L6 in SSp-bfd. Right, 3D-reconstructions of L2/3 and L6 targeted injection sites warped into the 3D-rendered space of SSp-bfd of the CCFv3 (Wang et al., 2020). Yellow colors indicate L2/3 targeted and purple colors indicate L6 targeted injection sites (n=2 mice for each condition). (I) Left, horizontal projection of VISp. Postsynaptic neurons labeled by both L2/3 and L6 targeted injections were warped to VISp of the CCFv3. Middle, fraction of postsynaptic neurons in VISp labeled by either L2/3 or L6 targeted injections to SSp-bfd. Right, Average fraction of postsynaptic neurons in different cortical layers (numbers) of VISp labeled by either L2/3 or L6 targeted injections to SSp-bfd. Numbers indicate cortical layers. Values are normalized to the maximum obtained after L6 or L2/3 targeted injections, respectively. (J) Density map of postsynaptic neurons in VISp (horizontal projection) of 5 individual mice. Border colors indicate the corresponding injection sites in VISp (see Figure 4F). (K) Horizontal projection to VISp. Crosses indicate the maxima of density of the 5 different populations of projections neuron. Colors correspond to the different injection sites in VISp. (L) Horizontal projection of VISp. Grey circles indicate postsynaptic neurons labeled by all 5 injection sites in VISp. Red arrows indicate the first and second principal components determining the direction of spatial dispersion of neuron locations. Black lines are section borders orientated vertically to the first principal component. (M) Average fraction of projection neurons was counted in all sections of VISp. (N, O) Average elevation and azimuth contour plots of VISp obtained by GCaMp6 fluorescence-based visuotopic imaging (Zhuang et al., 2017). Dashed black line: mean field sign borders of VISp. Solid black line: horizontal projection of the areal border of VISp from the CCFv3. Solid colored lines represent the horizontal meridian. Individual contours are 5° apart (modified from (Zhuang et al., 2017)).

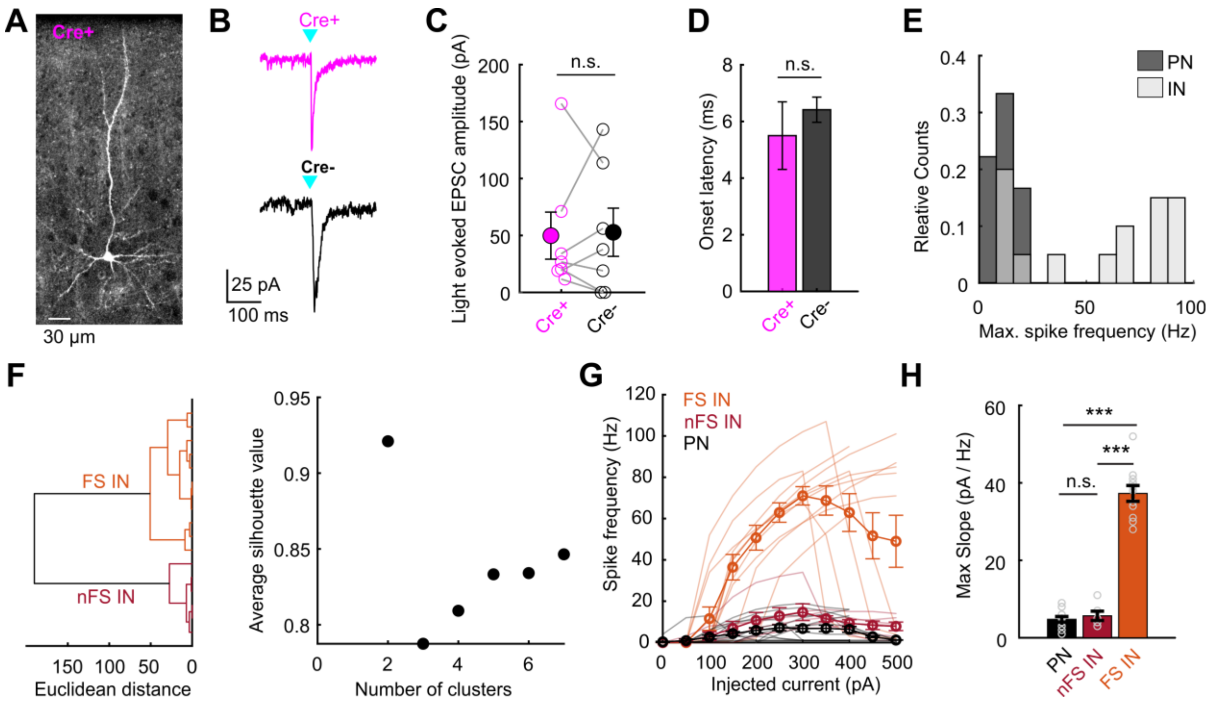


**Supplemental Figure 5: SSp-bfd mediated feedforward inhibition onto L2/3 neurons in V1.** (A) Anterogradely labelled L2/3 cell (Cre+) in VISp following AAV.hSyn.Cre injection in SSp-bfd of Ai14 mice. Scale bar: 30 µm. (B) Representative example of light-evoked EPSC in a Cre+ and a Cre- L2/3 cell. Scale bars: 25 pA, 100 ms. (C) Comparison of evoked peak EPSCs for Cre+ and Cre- cells. Mean (filled circles) and individual data points (empty circles) are displayed. Lines connect neighbouring cells (<100 µm apart; n=7 cells from 5 mice). Error bars are s.e.m. (p=0.87, Wilcoxon rank-sum). (D) Onset latency of light-evoked EPSCs for Cre+ and Cre- (n=6 and 13 cells from 6 mice). Error bars are s.e.m. (p=0.15, Wilcoxon rank-sum). (E) Distribution of maximal spike frequency for excitatory PNs and inhibitory interneurons (PN, IN; n=13 and 17 cells). Maximal spike frequency was obtained using current injections. (F) Left, Dendrogram of unsupervised hierarchical cluster analysis using Ward's method showing two major groups of interneurons based on their maximal spike frequency. Right, Average silhouette values plotted against cluster number. Note that the silhouette value is highest for two clusters. (G) Maximal spike frequency plotted against injected current amplitude for PNs, nFS INs and FS Ins (n=13, 6 and 11 cells). Mean (thick lines with circles) and individual data (thin lines) are displayed. (H) Maximal slopes for curves in (G) for PNs, nFS INs and FS INs. Mean and individual data points are displayed. Error bars are s.e.m. (p<0.001, Kruskal-Wallis).
